# Intra-DNA k-mer Conservation Patterns Encode Evolutionary Selection of Variants

**DOI:** 10.1101/2024.07.05.602168

**Authors:** Bernadette Mathew, Abhishek Halder, Nancy Jaiswal, Smruti Panda, Debjit Pramanik, Sreeram Chandra Murthy Peela, Abhishek Garg, Sadhana Tripathi, Swarnava Samanta, Prashant Gupta, Vandana Malhotra, Gaurav Ahuja, Debarka Sengupta

**Author notes:** Corresponding author*: { }. Equal contribution (Nancy Jaiswal, Smruti Panda). Equal contribution (Bernadette Mathew, Abhishek Halder).

## Abstract

Evolution shapes the structure and content of genomes, yet the contribution of local sequence composition to variant selection remains poorly understood. While traditional models emphasize protein function or cross-species conservation, we propose that intra-genomic patterns of oligonucleotide (k-mer) frequencies also reflect selective forces. To explore this, we developed **kGain score**, a metric that quantifies the frequency shift of a k-mer upon single-nucleotide substitution, using the surrounding genomic context as a baseline. We hypothesize that variants arising in high-kGain contexts are more likely to persist due to evolutionary favorability. We validated this hypothesis across multiple systems. In *E. coli* and *S. cerevisiae* long-term evolution experiments, we found that fixed, essential, and parallel mutations consistently show elevated kGain scores. This trend held in SARS-CoV-2 variants of concern and in an in-house antibiotic adaptation experiment, where a high-kGain *fusA* Y515N mutation conferred resistance and maintained fitness when overexpressed, demonstrating a causal link between kGain and adaptive potential. To enable cross-species generalization, we trained a transformer-based neural network regressor on LTEE-derived mutations to predict kGain from sequence alone. The model achieved high correlation in held-out in-domain data (Pearson *r* = 0.81) and accurately predicted kGain trends in out-of-domain data (Pearson *r* = 0.82), demonstrating that k-mer-based sequence constraints learned from one genome can be effectively transferred to others. Together, our results establish kGain as a biologically meaningful, scalable metric for probing within-genome sequence selection, offering a complementary lens to existing conservation-based frameworks for understanding evolutionary fitness and variant persistence.

## Introduction

A central goal in evolutionary genomics is to understand how genetic variation translates into phenotypic consequences and, ultimately, adaptive fitness across successive generations. To achieve this, computational methods for variant effect prediction (VEP) ^1^ have become critical tools for linking genotype to phenotype. Early VEP approaches, such as SIFT and PolyPhen, largely relied on evolutionary conservation signals extracted from cross-species alignments ^2^ to estimate the functional impact of mutations ^3^. These methods modelled the probabilities of reference versus alternate alleles by leveraging homologous protein sequences. More recently, large-scale deep learning frameworks, exemplified by ESM ^4^, Evo ^5^, and AlphaMissense ^6^, have advanced the field by capturing complex patterns of selection and functionality in protein-coding regions. Although these techniques have improved our ability to interpret coding variants, they often focus on interspecies conservation and may overlook subtle intra-genomic patterns of selection that govern the organization and evolution of the entire genome within a species.

Increasing evidence suggests that analyzing k-mer frequencies within a single genome offers a complementary window into how selection shapes genomic architecture ^7^. k-mer-based approaches, which quantify the occurrence of all possible short nucleotide sequences (k-mers), have proven valuable in dissecting genome composition and evolution. Additionally, differential k-mer frequencies within a single genome can reflect local selection pressures acting on various genomic elements, including transcription factor binding sites and regulatory motifs. k-mer analysis has thus been employed to illuminate how specific sequence elements are conserved within a genome, an idea that extends beyond the classical paradigm of protein-centric or cross-species conservation.

Such patterns of **within-genome conservation are** evident in microbial genomes, where selective pressures act rapidly, favoring certain nucleotide motifs essential for genomic stability/adaptation. For instance, bacteria often exhibit conserved k-mers within operons that enable synchronized gene expression, even in the absence of broader homology across species ^8^. Likewise, transposable elements, insertion sequences, and integrons maintain internal sequence motifs critical for their mobility and regulatory functions, suggesting that particular k-mers are preferentially retained for their functional utility within a given genomic context ^9,10^. In extremophilic archaea, duplicated stress-response genes often contain conserved k-mer motifs that enhance their tolerance to extreme temperatures or salinities, underscoring the potential adaptive significance of these patterns ^11^. Moreover, horizontal gene transfer, the primary driver of microbial evolution, frequently transfers regulatory elements or other k-mer motifs that seamlessly integrate into recipient genomes, affecting gene stability, expression, and evolutionary trajectories ^12,13^.

Together, these observations point to the possibility that certain k-mer motifs might be systematically favored during adaptive processes within the same genome, forming an underappreciated layer of selection that complements the traditional view of across-species sequence conservation. Despite the promise of these k-mer-based insights, the connection between specific k-mer motifs and their direct impact on evolutionary fitness remains underexplored. While k-mer-based methods have been successfully used in pangenomics and population genetics ^14,15^ revealing genetic diversity overlooked by single-reference-based approaches, a quantitative demonstration that k-mer frequency biases can drive or accompany adaptive changes has been lacking.

Here, we introduce a quantitative framework for evaluating whether k-mer frequencies within a genome are non-randomly selected over evolutionary time. **kGain score** is central to our approach, designed to capture the enrichment level of specific k-mers near single-nucleotide variants (SNVs). By examining multiple data sets, long-term evolution experiments (LTEE) in *E.coli* and *S.cerevisiae*, as well as natural evolutionary data of SARS-CoV-2, we provide evidence that variant selection correlates with systematic shifts in genome-wide k-mer frequencies. We also performed a single-colony bottleneck passage experiment in *E. coli* under sublethal antibiotic pressure, revealing that k-mer frequency dynamics are not merely observational artifacts but can have tangible fitness consequences under controlled selection regimes.

Our results yield four key findings. First, we show that the median kGain score of given variants in *E. coli* and *S. cerevisiae* populations tends to increase over the course of evolution, suggesting a bias toward certain k-mer enrichments ^16^. Second, essential genes preferentially accumulate variants with higher kGain scores, indicating a functional link between k-mer selection and vital cellular processes. Third, fixed mutations, those that persist over multiple generations, consistently exhibit higher kGain values, underscoring the evolutionary advantage conferred by these motifs. Finally, imposing sublethal antibiotic pressure in a single-cell bottleneck experiment amplifies selection for high-kGain variants, tying k-mer frequency biases directly to adaptive stress responses.

Collectively, these insights propose an expanded view of molecular evolution, one that considers the potential role of k-mer frequencies within the same genome, intra **genome conservation**, as a critical, yet largely overlooked, layer of selection. By tracking k-mer dynamics through the kGain metric, we establish a new avenue for identifying variants that may be key to adaptation. As genome sequencing efforts broaden across diverse organisms and ecological contexts, this framework holds promise for enhancing our understanding of how genomic architecture evolves under selective pressures, complementing the well-established narratives of cross-species conservation and protein-focused evolutionary dynamics.

## Results

### k-mer frequency patterns within and between genomes

k-mers are short nucleotide sequences commonly used in bioinformatics for similarity searches, sequence alignment, and homology detection ^17,18^. To examine whether k-mers experience selective pressures, we explored their non-uniform distribution within individual genomes, followed by an investigation into the underlying principles behind such bias. To visualise k-mer frequencies, we generated Frequency Chaos Game Representations (FCGR) using Python (kaos) package ^19^.

In our analysis, we represented k-mer frequencies using the Frequency Chaos Game Representation (FCGR), in which each k-mer is assigned to a specific coordinate in a two-dimensional matrix (**Fig. 1a**), thereby encoding global sequence composition. As a benchmark, we computed absolute frequency differences for randomly paired k-mers sampled from *E. coli* genomes and found that pairs differing by a single nucleotide exhibited significantly smaller disparities than random pairs, a pattern that persisted up to Hamming distances of three (**Fig. 1b**). This observation reflects the FCGR’s recursive partitioning, which clusters k-mers sharing a common prefix into neighboring matrix regions and thus preserves sequence similarity in spatial proximity (^20^). When applied to diverse genomes, the FCGR revealed fractal-like motifs in mammalian sequences, for example, human (*Homo sapiens*; **Fig. 1c**), gorilla, and mouse (*Mus musculus*; **Supplemental Fig. S1a–b**), whereas more discrete, non-fractal patterns emerged in zebrafish (*Danio rerio*; **Supplemental Fig. S1c**) and yeast (*Saccharomyces cerevisiae*; **Fig. 1d**) ^21^.

**Figure 1.**
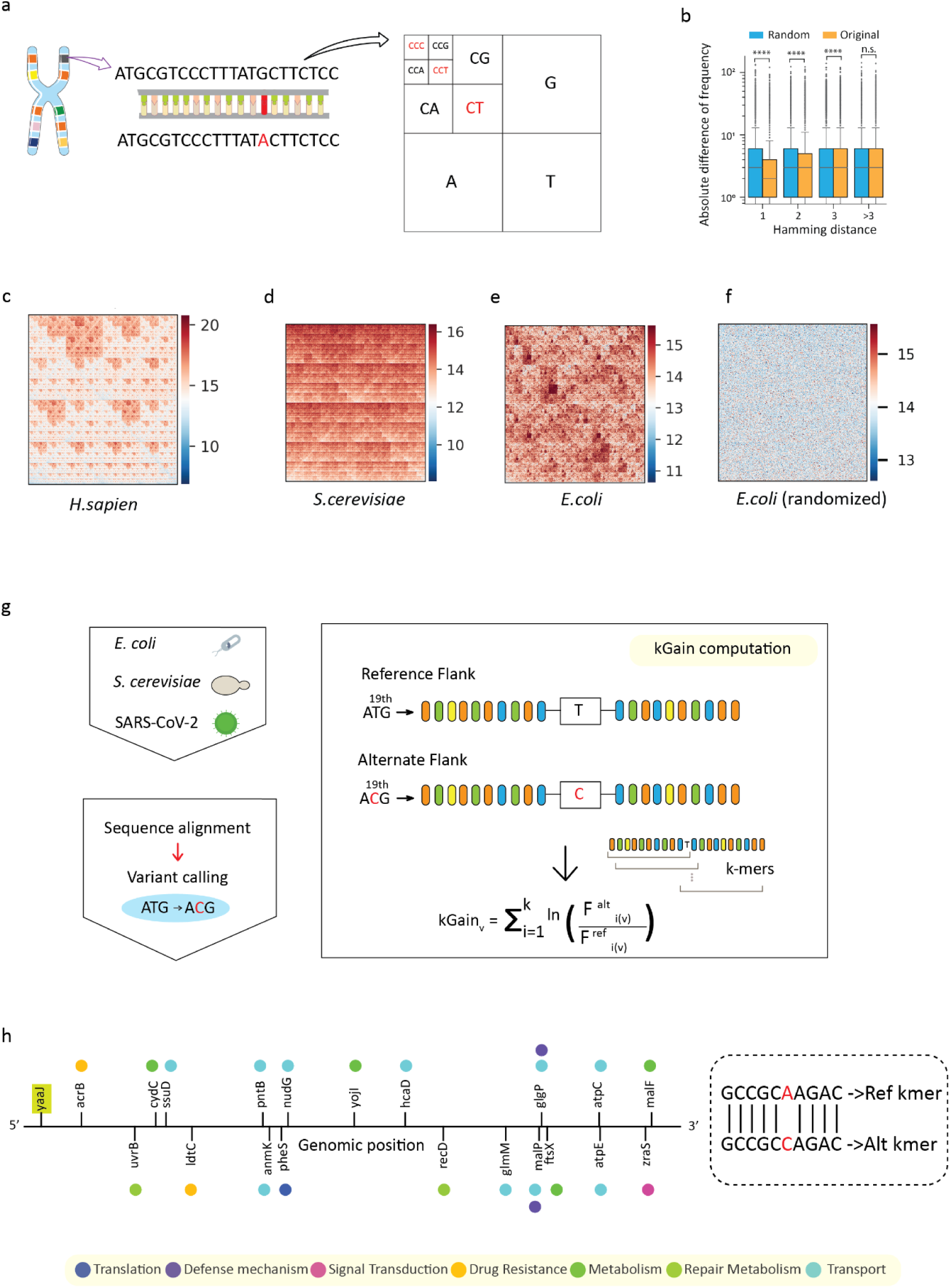
Exploring the fractal pattern of DNA across various organisms and examining its non-random nature. **(a)** The genomic DNA is fragmented into oligonucleotides of a defined length *k*, which are subsequently mapped to the FCGR matrix of the reference genome. **(b)** Boxplot showing the absolute difference (log scale) in frequency distribution between two randomly selected *k*-mers and one randomly selected *k*-mer with its counterpart altered by n Hamming distances (original altered) (*P*-values of <1e-323 for Hamming distances 1 and 2, 1.56e-89 for distance 3, and 0.09 for distances greater than 3). **(c-e)** Heatmaps illustrate normalised frequency values (in negative logarithmic scale) to reduce the range, which demonstrates that various organisms exhibit distinct patterns of *k*-mer abundance within their DNA sequences. **(f)** To evaluate the significance of these fractal patterns, we utilised a heatmap of genomic DNA from *E. coli*, comparing it to a randomised counterpart, thereby elucidating discernible differences in pattern preservation. **(g)** kGain score computation process: For each variant, two 21-mer/13-mer sequences are generated, one with the reference allele and one with the alternate allele at the central position. k-mers (k = 10) are generated using a rolling window method for both reference and variant sequences, resulting in sets of k-mers. The kGain score for each variant is calculated by summing the natural log of the fold change between the genomic frequencies of the k-mer containing the alternate allele and the reference allele across all k-mers. **(h)** The schematic illustrates the distribution of the alternate k-mer (resulting from an A→C substitution) across various genomic loci in *E. coli*. The *yaaJ* gene, highlighted in green, serves as the reference k-mer source. The alternate k-mer is mapped across multiple genes involved in diverse biological functions, including translation, metabolism, drug resistance, transport, defence mechanisms, and signal transduction. The functional classification of each gene is denoted by colored circles. The boxed region on the right depicts the reference (Ref) and alternate (Alt) k-mer sequences, highlighting the nucleotide substitution in red.

Notably, these FCGR k-mer frequency-based correlations preserve phylogenetic relationships across species ^22,23^ (**Supplemental Fig. S1d**). Although k-mer frequency conservation across closely related species is well captured by phylogenetic trees, the intra-genomic rationale for differential k-mer frequencies remains largely unexplored^15^. To explore whether genome-specific k-mer patterns are selectively maintained, we generated randomised *E. coli* genomes while preserving overall nucleotide ratios. In contrast to the original *E. coli* sequence, these simulated genomes displayed no discernible fractal pattern in their FCGR heatmaps (**Fig. 1e–f**). Could these reflected fractal patterns arise because near-identical k-mers exhibit comparable frequencies across the entire genome, a feature that is biologically significant for selection? To explore this possibility, we investigated whether similar near-identical k-mers within an *E. coli* genome share similar frequencies ^24^. These findings suggest the systematic conservation of k-mers within an individual genome.

In our pursuit to investigate whether such differential k-mer frequencies are linked to evolution, we hypothesized that SNVs under selection might replace low-frequency k-mers with high-frequency ones. To test this, we developed a simple score, kGain, which quantifies the relative abundance of k-mers associated with alternative alleles compared to reference alleles (**Fig. 1g**). For further information on kGain computation, please refer to the **Methods** section. For all our analysis, the value of k was set to 10 (**Supplemental Fig. S1e-f**, **Supplementary Note 1**). To illustrate how k-mers are captured within a genome, we used *yaaJ* A7764C as an example (**Fig. 1h**). In the following sections, we leverage the kGain score to explore the link between within-genome differential k-mer frequencies and selection.

### k-mer Enrichment Patterns Reflect Selective Pressures Across Evolutionary Contexts

To evaluate whether local sequence composition influences the retention of genetic variants, we analyzed k-mer frequency dynamics across diverse evolutionary landscapes, including laboratory evolution, natural viral adaptation, and controlled antibiotic stress. We first examined high-resolution longitudinal whole-genome sequencing data from the long-term evolution experiments (LTEE) in *Escherichia coli* and *Saccharomyces cerevisiae*, which capture the stepwise accumulation of mutations across tens of thousands of generations under defined, nutrient-limited conditions. These systems provided a robust framework for observing how adaptive trajectories unfold over time. To extend our analysis into natural evolution, we analyzed global genomic surveillance data from SARS-CoV-2, leveraging the emergence of major viral variants during the COVID-19 pandemic as a model for rapid, large-scale adaptation driven by host immunity, transmission dynamics, and medical interventions. Finally, to assess how acute selective pressures shape k-mer profiles in real time, we performed a de novo single-colony bottleneck evolution experiment in *E. coli* exposed to incrementally increasing sublethal kanamycin concentrations. This controlled setup allowed us to monitor evolutionary responses under antibiotic stress, directly linking mutational dynamics to survival outcomes (**Fig. 2a–d**). Together, these datasets provide a unified framework for assessing whether, and how k-mer frequency changes accompany evolutionary selection across timescales and ecological settings.

**Figure 2.**
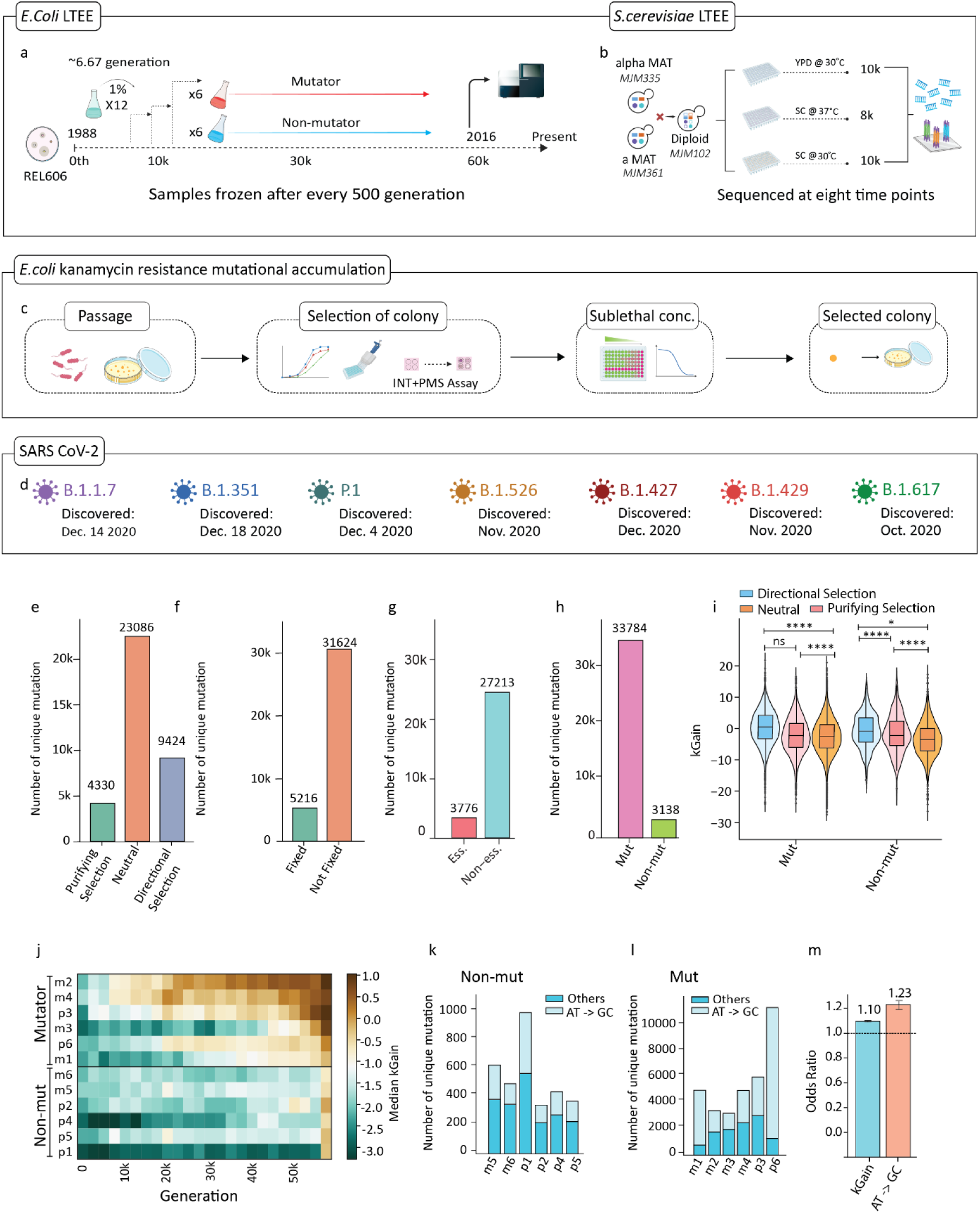
Evolutionary dynamics in *E. coli* LTEE dataset. **(a)** A visual depiction of the LTEE involving *Escherichia coli B* strain REL606. Initiated in 1988 by Lenski et al., this experiment has spanned over 30 years and includes 12 replicate populations labelled ara-1 to ara-6 (m1 to m6) and ara+1 to ara+6 (p1 to p6). These populations were propagated for 60,000 generations (1998-2016) and subjected to sequencing at intervals of 500 generations. **(b)** Schematic overview of the *Saccharomyces cerevisiae* LTEE: The experiment began with alpha MAT (MJM335) and a MAT (MJM361) strains, which were crossed to generate a diploid (MJM102). These populations were evolved in three different environments: YPD at 30°C for 10,000 generations, SC at 37°C for 8,000 generations, and SC at 30°C for 10,000 generations. Samples were sequenced at eight time points throughout the experiment. **(c)** Schematic representation of the experimental workflow for bacterial adaptation under sublethal antibiotic pressure. The process begins with the passage of bacterial populations, followed by colony selection based on growth characteristics measured using the growth curve or INT+PMS assay. After selecting the colony, sublethal concentrations of antibiotics are determined, and the selected colony is grown in the following passage. **(d)** Here is an example of the key variants of SARS-CoV-2 tracked in this study, showing the lineage names, color-coded symbols, and dates of first discovery. **(e)** A bar plot representing unique mutations classified by selective regime, purifying selection (green), drift (orange), and directional selection (blue), based on temporal allele frequency trajectories. **(f)** A bar plot showing the number of unique mutations stratified by fixation status, with fixed mutations (green) and not fixed mutations (orange) identified across all populations. **(g)** A bar plot illustrating the distribution of unique mutations among essential genes (red) and non-essential genes (cyan). **(h)** A bar plot depicting the number of unique mutations in mutator (pink) and non-mutator (green) gene sets. **(i)** Violin and box plots showing the distribution of kGain scores for mutations under directional selection (blue), drift (orange), and purifying selection (red) in both mutator (*P*-value <1e-323 and effect size = 7.97e-01 for directional selection vs neutral in mutator, *P*-value = 1.63e-134 and effect size = 7.41e-01 for directional selection vs purifying selection in mutator, *P*-value = 9.95e-01 and effect size = -4.89e-02 for neutral vs purifying selection in mutator) and non-mutator populations (*P*-value = 2.97e-17 and effect size = 7.57e-01 for directional selection vs neutral in non mutator, *P*-value = 3.03e-02 and effect size = 3.75e-01 for directional selection vs purifying selection in non mutator, *P*-value = 5.56e-06 and effect size = 3.70e-01 for purifying selection vs neutral in non mutator). **(j)** Heatmap showing the median kGain values across generations for individual populations, stratified by mutator and non-mutator. (**k**) Number of unique mutations in non-mutator populations, separated by mutation type (A/T→G/C vs. Others). (**l**) Number of unique mutations in mutator populations, separated by mutation type. (**m**) Odds ratio comparing the enrichment of kGain and A/T→G/C substitutions. Error bars indicate 95% confidence intervals. [**Note**: The *P*-value cutoff for all the plots is 0.05. *, **, ***, and **** refers to p-values <0.05, <0.01, <0.001, and <0.0001, respectively.]

We evaluated several critical dimensions to interpret the significance of kGain scores, including the classification of mutations as purifying, directional, or neutral ^25^; the fixation status of variants (fixed vs. not-fixed); the distinction between essential and non-essential gene; and the genomic context of mutator versus non-mutator lineages (**Fig. 2e-h**). As a first step, we applied kGain to reanalyze genomic data from the landmark *E. coli* LTEE initiated by Lenski and colleagues ^26,27^. Guided by established population genetic theory (e.g., neutral theory 1968), we categorized mutations identified in the LTEE into three classes, positively selected (directional selection), negatively selected (purifying selection), and neutral, based on the trajectories of their allele frequencies over time^26,27,28#x2013;30^. Mutations were classified by modeling allele-frequency trajectories on the log-odds scale and testing the per-generation change in frequency. Mutations with a significantly positive slope (β_1_ > 0, p < 0.05) were classified as positively selected, those with a significantly negative slope (β_1_ < 0, p < 0.05) as negatively selected, and mutations without a significant trend (p > 0.05) were considered neutral **(Supplementary Note 2)** . We investigated certain aspects like the directional selection, purifying selection and genetic drift (neutral) along with the AT to GC conversion. We noticed directional selection was associated with higher kGain scores compared to both neutral and purifying selection (**Fig. 2i**). We further analyzed kGain scores in mutator and non-mutator backgrounds and observed a gradual increase in the abundance of alternate k-mers over time in both groups (**Fig. 2j**), reflecting the progressive accumulation of high-frequency alternate k-mers mutations under differing mutation rates.

We examined the count of A/T to G/C conversions in both non-mutator (**Fig. 2k**) and mutator (**Fig. 2l**). As shown in **Fig. 2m**, the odds of a mutation being classified as beneficial were 23% higher for A/T to G/C substitutions than for other base changes (odds ratio = 1.23). High-kGain loci also exhibited increased odds of being beneficial (OR = 1.10), indicating that both the mutation class and kGain metric are associated with the predicted functional impact of mutations (**See Methods**). Next, we examined whether generation-wise kGain scores exhibit a systematic temporal pattern when calculated using a population-mutated evolved genome (constructed by introducing mutations observed in the final generation into the wild-type reference). We observed a clear increasing trend in k-mer enrichment over time, with the rate of increase gradually plateauing in later generations, a biologically meaningful signature consistent with mutation accumulation dynamics (**Fig. 3a**). To evaluate whether this pattern could arise by chance, we performed a permutation test, which yielded a Monte Carlo P = 1×10⁻⁴ (CI = 0.768), indicating that the observed trend is highly unlikely to occur randomly and underscores kGain’s robustness in capturing temporal shifts in sequence composition during evolution (**Supplementary Note 3**). To benchmark our findings against two state-of-the-art cross-species protein/DNA sequence foundation models, we evaluated ESM1b (650 million parameters) ^4^ and Evo (7 billion parameters) ^5^,as reference standards. These models output the probability of observing an input given an amino acid (ESM1b) or nucleotide (Evo) sequence across species. Notably, kGain is fundamentally different from these two since it solely focuses on k-mer frequencies within a single reference genome. Log Likelihood Ratio (LLR) score provided by ESM1b (**Fig. 3b**) (C.I.: 0.64, Monte Carlo *P*-value: 1e-4) showed the best concordance, followed by kGain **(Supplementary Note 4)**. No significant association was detected with Evo (**Fig. 3c**) (C.I. : -0.40, Monte Carlo *P*-value: 1) . We observed these trends consistently across individual populations as well. If kGain effectively captures selection-driven shifts in genomic composition, we sought to validate this by testing it against a well-characterized evolutionary transition reported by Blount and colleagues. In the *E. coli* LTEE, population m3 exhibited a dramatic phenotypic change around generation 33,000, marked by a sudden increase in turbidity due to the evolution of citrate utilization under aerobic conditions, a major adaptive innovation. When we examined the corresponding kGain trajectory, we observed a distinct elevation in median kGain values precisely at this transition point. This alignment suggests that kGain can sensitively register genome-wide shifts in selection pressure during key adaptive events^31^ (**Supplemental Fig. S1i**). Based on our observation that kGain may reflect the selective retention of beneficial mutations, we tested whether a simple linear model, using short k-mer frequencies, could predict kGain. Although this approach explained part of the variation (R2 = 0.49, RMSE = 3.80, Pearson correlation = 0.70 on the test set), it fell short of precise prediction (**Supplemental Fig. S2a-g**), as detailed in **Supplemental Note 5.**

**Figure 3.**
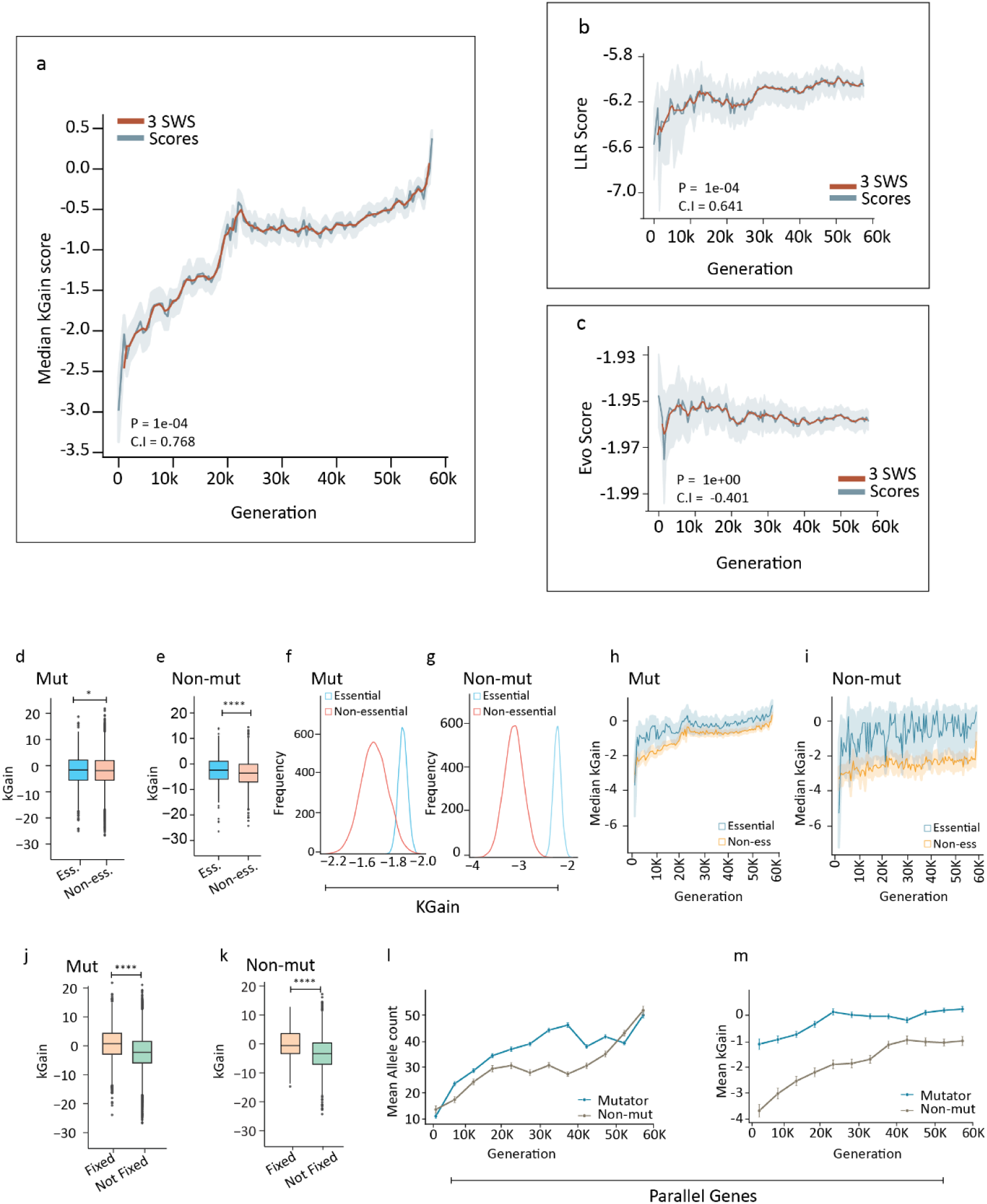
Beneficial mutational spectrum shift. **(a)** Median kGain across generations, with 95% confidence intervals shown as shaded areas. **(b)** Temporal trends of median LLR score over generations. **(c)** Median Evo score over generations. **(d)** Box plot of kGain scores in mutator populations comparing essential and non-essential genes (*P*-value = 2.40e-02 and effect size = 8.00e-02). **(e)** Box plot depicting kGain scores in non-mutator populations comparing essential and non-essential genes (*P*-value = 5.14e-05 and effect size = 3.05e-01). **(f)** Bootstrapped distributions of mean evolved kGain scores for essential and non-essential gene mutations in mutator populations, generated using 10,000 iterations and subsampling 90% of the minimum group size per iteration. **(g)** Bootstrapped distributions of mean evolved kGain scores for essential and non-essential gene in non mutator populations, generated using 10,000 iterations and subsampling 90% of the minimum group size per iteration. **(h)** Median kGain score over generations for essential and non-essential genes in mutator populations. **(i)** Median kGain score over generations for essential and non-essential genes in non-mutator populations. **(j)** Box plot of kGain scores in mutator populations comparing fixed and not fixed mutations (P-value = 1.04e-242 and effect size = 7.94e-01). **(k)** Box plot of kGain scores in non-mutator populations comparing fixed and not fixed mutations (*P*-value = 7.95e-14 and effect size = 7.69e-01). **(l)** Line plot of mean allele count per generation for mutator and non-mutator in parallel genes. **(m)** Line plot of generation wise mean kGain for mutator and non-mutator in parallel genes. [**Note**: The p-value cutoff for all the plots is 0.05. *, **, ***, and **** refers to p-values <0.05, <0.01, <0.001, and <0.0001, respectively.]

### Quantitative Dynamics of kGain Enrichment in Essential Genes: Fixed, Non-Fixed, and Parallel Mutation Profiles

Understandably, essential genes are generally resilient to mutations, as mutation can impact the function and therefore the organism’s survival or fitness. Intuitively, if kGain scores truly indicate evolutionary advantage in essential genes, we would observe a bias towards variants with higher kGain scores. This is because essential gene mutations with poor kGain scores would often have a deleterious impact on the organism, thereby eliminating populations carrying the variant allele. We sourced essential/non-essential categorization from the study by Gerdes et al. ^32^. Aligned to our hypothesis, we found elevated median kGain scores for single nucleotide substitutions associated with the essential genes, as compared to those associated with the non-essential ones (**Fig. 3d,e**). Bootstrapped distributions of mean kGain scores for essential and non-essential mutations in mutator and non-mutator populations, generated using 10,000 iterations with 90% subsampling of the minimum group size per iteration, revealed a similar pattern of enrichment (**Fig. 3f,g**). Longitudinal analysis indicated that generation wise median kGain scores for essential genes remained consistently above those of non-essential genes throughout the experiment, reflecting the stronger constraint and purifying selection operating on essential loci (**Fig. 3h,i**). Unlike kGain, LLR scores did not exhibit a comparable separation between essential and non-essential genes, suggesting that LLR is less sensitive to such functional constraints.**(Supplemental Fig. S3a-c)**.

We compared kGain scores between fixed and not fixed mutations in both mutator and non-mutator backgrounds. Fixed mutations were defined as those with allele frequencies greater than or equal to 0.95 at the last two observed time points of the experiment (**Supplementary Note 6**). We observed that fixed mutations exhibited significantly higher kGain scores than those not fixed at the endpoint in both mutator and non-mutator populations (**Fig. 3j,k and Supplemental Fig. S3d,e**). This pattern suggests that mutations which reach fixation are typically under stronger selection, consistent with the expected outcomes of adaptive evolution.

To assess whether loci repeatedly mutated across independent lineages, the so-called *parallel genes*, harbor signatures of adaptive selection, we analyzed their kGain trajectories in comparison to non-parallel genes, as defined by ^30^. Across the LTEE, parallel genes, recurrently mutated across independent populations, consistently showed elevated median kGain scores, with this enrichment being especially prominent in mutator lineages compared to non mutator. As an additional control, we examined allele count trajectories, which revealed similar trends (**Fig. 3l,m**).

We binned mutations into 5,000-generation intervals and calculated median allele frequencies for each bin, stratified by mutator status (mutator vs. non-mutator) and parallelism status (parallel vs. non-parallel genes). Median allele frequency trajectories were visualized using smoothed line plots and overlaid scatter points representing bin-wise medians. To test whether parallel gene mutations exhibited consistently higher allele frequencies compared to non-parallel gene mutations within each generation bin, we performed a one-sided Mann–Whitney U test separately for mutator and non-mutator populations. Parallel gene mutations in mutator populations rapidly rise to high allele frequencies than non-parallel genes, with significance across most generation bins. Non-mutator populations also favor parallel mutations but at lower frequencies, while non-parallel genes remain largely neutral (**Fig. 4a**). A similar pattern was observed for median kGain, with parallel gene mutations, showing consistently stronger positive selection signals compared to non-parallel genes (**Fig. 4b**).

**Figure 4.**
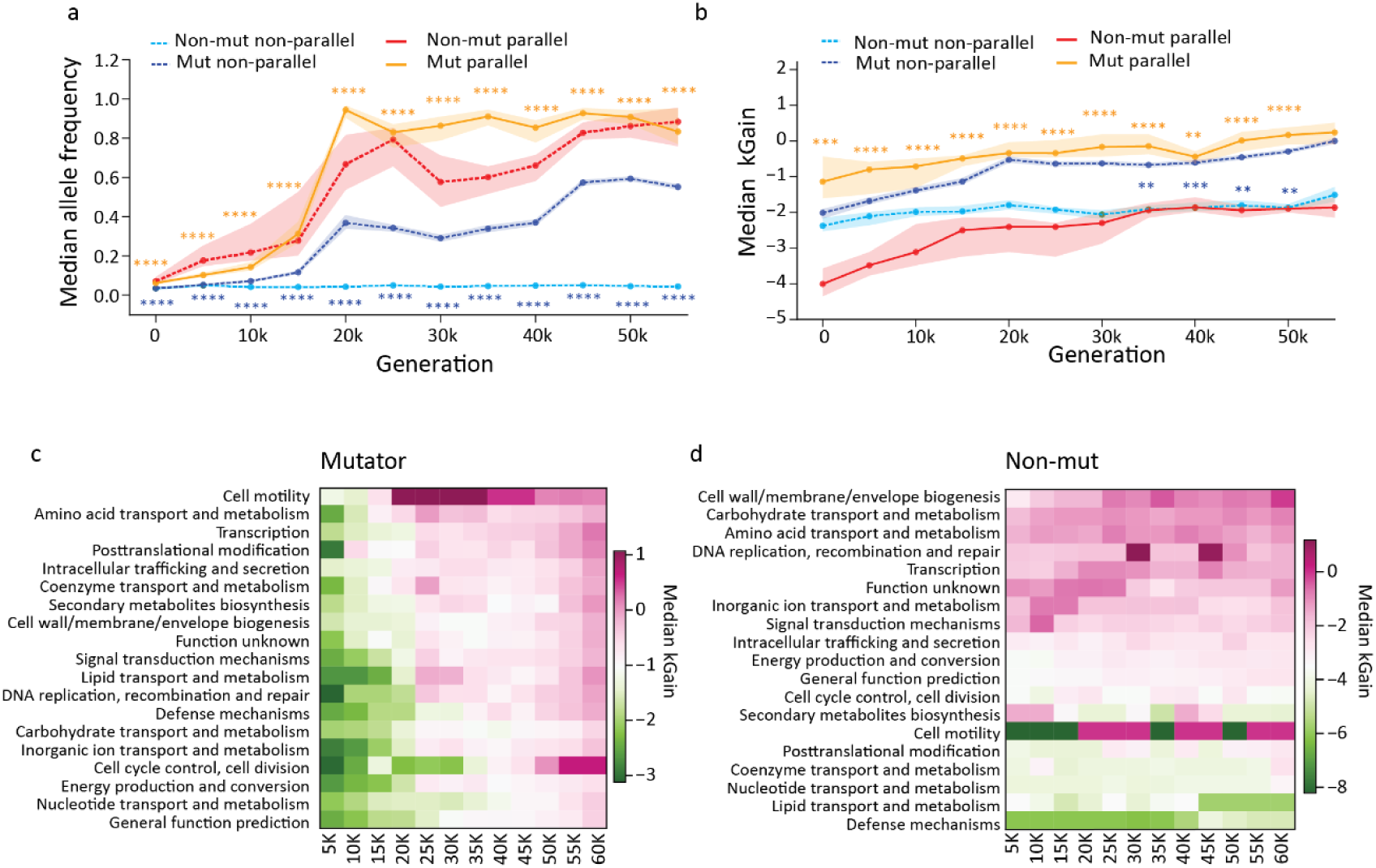
Temporal dynamics of allele frequency, kGain scores, and functional signatures in parallel and non-parallel genes in mutator and non-mutator backgrounds. **(a)** Temporal trends of median allele frequency in parallel and non-parallel genes across mutator and non-mutator populations. **(b)** Median kGain scores over generations for parallel and non-parallel genes in mutator and non-mutator. Shaded regions represent 95% confidence intervals, and statistical significance between groups at each time point is indicated by asterisks. **(c)** Heatmap of median kGain scores for major COG functional categories over generation in mutator populations. **(d)** Heatmap of median kGain scores for major COG functional categories over time in non-mutator populations. [**Note**: The p-value cutoff for all the plots is 0.05. *, **, ***, and **** refers to p-values <0.05, <0.01, <0.001, and <0.0001, respectively.]

We then compared kGain distributions between coding and non-coding regions, focusing on essential versus non-essential genes. Essential genes consistently harbored variants with higher kGain scores, a trend most evident in non-mutator populations. This pattern implies that under lower mutation supply, selection more stringently filters variants, favoring those in high-kGain contexts that preserve core biological functions. Non-mutators, therefore, may be more efficient in retaining beneficial or tolerable mutations in essential coding regions while mutators, with broader exploration, show more diffuse kGain distributions **(Supplemental Fig. S3f–k)**. Together, these findings highlight how kGain reflects both evolutionary constraint and adaptive prioritization across functionally important genomic regions.

### Module-Specific kGain Dynamics in Mutator vs. Non-Mutator Lineages

In mutator populations (**Fig. 4c**), early generations (5K–20K) show broadly negative to neutral median kGain across nearly all functional modules, consistent with initial purifying selection or drift. From ∼30K generations onward, however, the majority of categories transition to positive kGain (pink/magenta) by 60K. The most pronounced enrichments occur in cell motility, amino acid transport & metabolism, transcription, posttranslational modification, and intracellular trafficking & secretion, indicating these systems become focal points of positive selection as adaptation unfolds. A handful of core functions, such as general function prediction and nucleotide transport & metabolism, remain closer to zero or slightly negative (light green), suggesting ongoing constraint or balanced turnover.

By contrast, non-mutator lineages (**Fig. 4d**) exhibit a largely static kGain landscape. Across all timepoints, most modules, including cell wall/membrane biogenesis, carbohydrate transport, DNA replication & repair, and energy production & conversion—remain neutral to mildly beneficial, with only transient (**Supplemental Fig. S4**). Even by 60K, non-mutators show little sustained positive kGain, underscoring that without an elevated mutation supply, adaptive shifts in k-mer frequencies are both weaker and less widespread.

### Intra-species generalization of kGain predicts mutational effects with a attention model

In biological systems, the effect of a mutation depends not only on the alternate allele but also on its neighbouring sequence, a key concept in modern evolutionary modeling. Traditionally, approaches such as ESM rely on inter-species conservation to predict mutational effects. Here, we investigated whether kGain can also capture mutational effects within a species by quantifying the influence of local sequence context. To test this hypothesis, we trained an attention-based neural network (see **Methods**) to predict kGain from sequence windows centered on the mutation site in the reference genome of *Escherichia coli B* strain REL606. We then evaluated whether the model could predict mutational effects, as represented by kGain scores, in other *E. coli* strains. Standard one-hot encoding for SNP regression often fails to encode the exact position or nature of the allele change within the sequence. To address this, we introduced a dual-encoded sequence representation (see **Methods**), which embeds both the reference flanking region and the alternate allele, providing the model with complete information about each mutation event (**Fig. 5a**). This encoding served as input to the transformer-based attention model. To further improve learning, we developed a custom loss function (see **Methods**) inspired by focal loss, emphasizing samples with higher prediction errors and encouraging the model to better fit challenging cases compared to standard loss functions.

**Figure 5.**
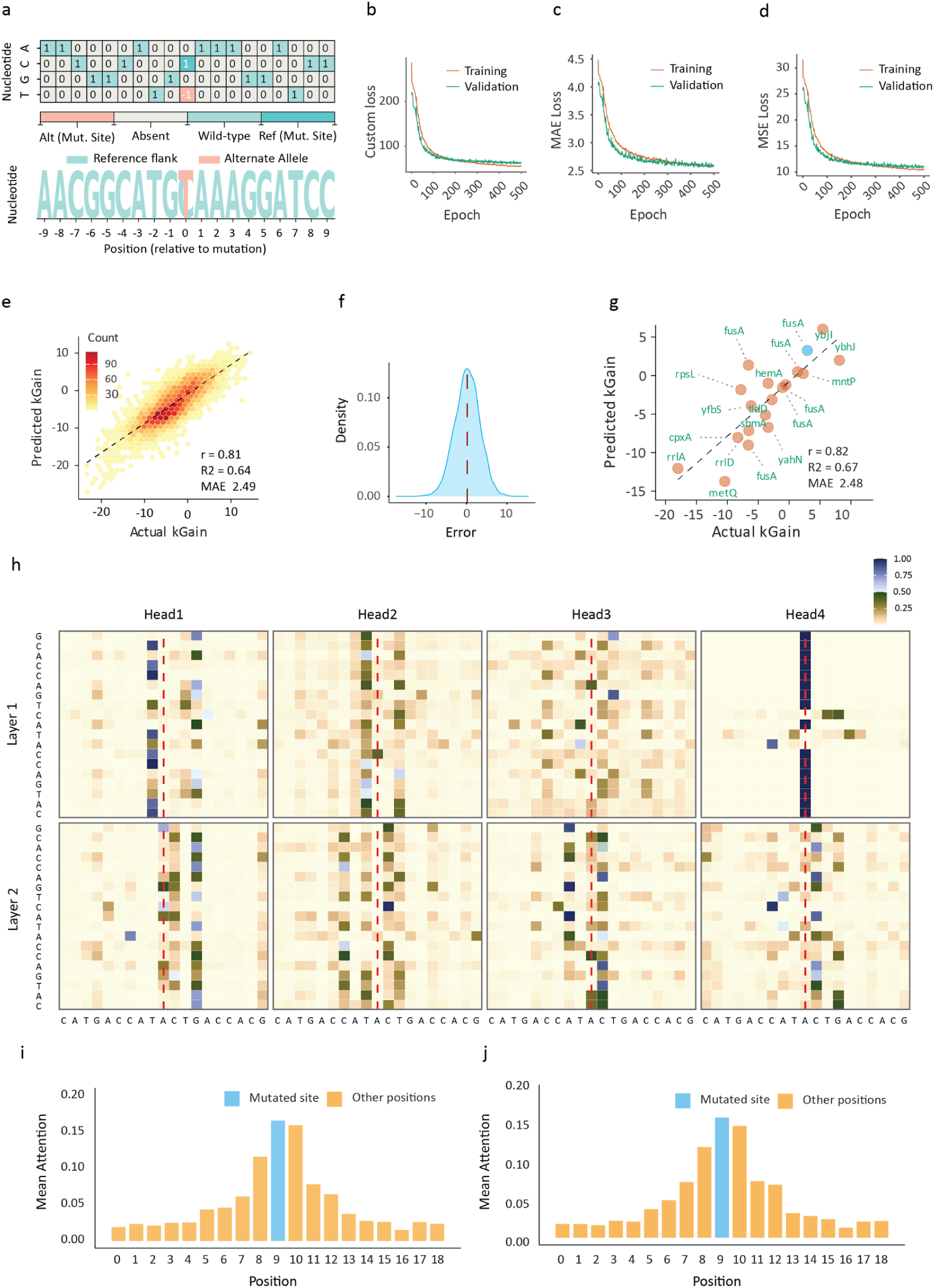
Performance and interpretation of the kGain prediction model. **(a)** Encoding strategy for representing SNP mutations: the panel illustrates the custom embedding approach used to encode single-nucleotide polymorphisms (SNPs) for the prediction model. The upper matrix shows the one-hot encoded states for each nucleotide (A, C, G, T) at positions flanking the mutation site, as well as at the mutation site itself. The lower sequence logo visualizes the nucleotide context around the mutation, highlighting the reference sequence, the alternate allele at the mutation site, and the reference flanking positions. This strategy captures both the local sequence and the specific nucleotide change introduced by each SNP. **(b)** Training and validation loss curves (c) MAE loss curve Training and validation **(d)** MSE loss curve Training and validation **(e)** In-domain performance. Predicted versus observed kGain scores for the held-out dataset, colored by point density, show strong agreement (Pearson correlation r = 0.81). **(f)** Distribution of prediction errors for the held-out LTEE data, confirming approximate normality with zero mean. **(g)** Out-of-domain validation. Predicted versus observed kGain scores for mutations from an independent E. coli strain (*E. coli MG1655*) demonstrate robust generalization of the model (Pearson correlation r = 0.82). **(h)** Attention map for the Y515N mutation in *fusA* in *E. coli* MG1655, illustrating the model’s focus on the mutation site and neighboring positions. **(i)** Average attention map across the sequence window for multiple samples from *E. coli* MG1655 in the in-house experiment. **(j)** Average attention map across the sequence window for held-out samples of *E. coli* B strain REL606 from the LTEE experiment.

For robust evaluation, we reserved 20% (6,879 samples) of the LTEE mutation data as an independent test set, using the remaining 80% (27,512 samples) for training and validation with an 80/20 split per epoch. The held-out set allowed us to rigorously assess in-domain performance. Both training and validation losses decreased steadily over 500 epochs (**Fig. 5b**), without evidence of overfitting. Additionally, we observed that optimizing the custom loss led to concurrent reductions in both mean absolute error and mean square error (**Fig. 5c-d**), further supporting the model’s generalizability. On the held-out test set, the model achieved a Pearson correlation of 0.81, ℝ^2^ of 0.64, and MAE of 2.49 between actual and predicted kGain scores. While the MAE was moderate, the high correlation demonstrates that the model effectively learns the ranking of mutational effects, even if the exact values are not perfectly predicted (**Fig. 5e**). The kGain error distribution in the LTEE dataset followed a Gaussian profile with a mean near zero and standard deviation of 3.14 (**Fig. 5f**). To test robustness to out-of-domain data, we evaluated the model trained solely on *E. coli B strain REL606*, on mutations from the *E. coli MG1655* strain (see “***Bacterial adaptation under antibiotic pressure validates kGain scores***” below), without any fine-tuning. In our in-house experiment, we found that the Y515N mutation in *fusA* was strongly selected, as its high kGain score predicted both robust kanamycin resistance and maintained growth fitness, making it an optimal adaptive mutation compared to other lower kGain variants. The attention model predicted a kGain value of 3.26 for this mutation, closely matching the actual value and demonstrating the model’s ability to generalize to other strains. When evaluated on the entire in-house dataset (out-of-domain), the model maintained strong performance (Pearson correlation = 0.82, ℝ^2^ = 0.67, MAE = 2.48; **Fig. 5g**), indicating that the kGain framework can generalize mutational effects across strains within a species. To further interpret the model’s predictions, we visualized attention maps (**Fig. 5h**) for the Y515N mutation in *fusA*. The maps revealed that the model focuses most strongly on the mutation site while also considering neighboring bases, consistent with biological principles. This pattern highlights how the dual-encoded sequence representation enables the model to explicitly recognize and interpret the mutation’s precise location. To determine whether this attention pattern was unique to one sample or represented a broader trend, we analyzed average attention maps across the sequence window for multiple samples. We found that all positions contributed to kGain prediction, with the greatest importance assigned to nucleotides near the mutation site. This influence decreased exponentially with distance from the mutation, in line with the rolling window approach for kGain calculation, where central nucleotides appear in more windows and thus have a larger impact on kGain. However, even distant positions contributed to the prediction, as reflected in the attention maps for both *E. coli B strain REL606* and *E. coli MG1655* (**Fig. 5i-j**). These findings underscore the importance of both local and broader sequence context in determining mutational effects.

### Bacterial adaptation under antibiotic pressure validates kGain scores

An in-house mutational-accumulation experiment under antibiotic pressure was performed to evaluate the persistence of k-mer based selection biases during adaptation to kanamycin. Wild-type *E. coli* K-12 MG1655 lineages were subjected to five successive serial passages in sublethal, incrementally increased kanamycin concentrations (0.006 → 0.1 mg·mL^-1^) using a single-colony bottleneck regime. One lineage (D) was maintained on nutrient-rich Luria–Bertani agar, while three parallel replicates (R1, R2, R3) were propagated in minimal M9 medium. Replicate R2 failed to re-establish growth after the second passage and was subsequently excluded. At each transfer, the colony capable of surviving a 2–4× increase in kanamycin concentration served as the inoculum for the next passage.

By the end of passage five, replicate R1 exhibited no detectable growth and was terminated. Growth kinetics of all the endpoint clones were then quantified both in the presence (D: 0.08 mg·mL^-1^; R1: 0.20 mg·mL^-1^; R3: 0.10 mg·mL^-1^) and absence of kanamycin in LB media.

Population D displayed growth indistinguishable from wild type under both conditions, whereas R1 and R3 exhibited significantly impaired proliferation even without antibiotic challenge (**Fig. 6a; Supplemental Fig. S5a**).

**Figure 6.**
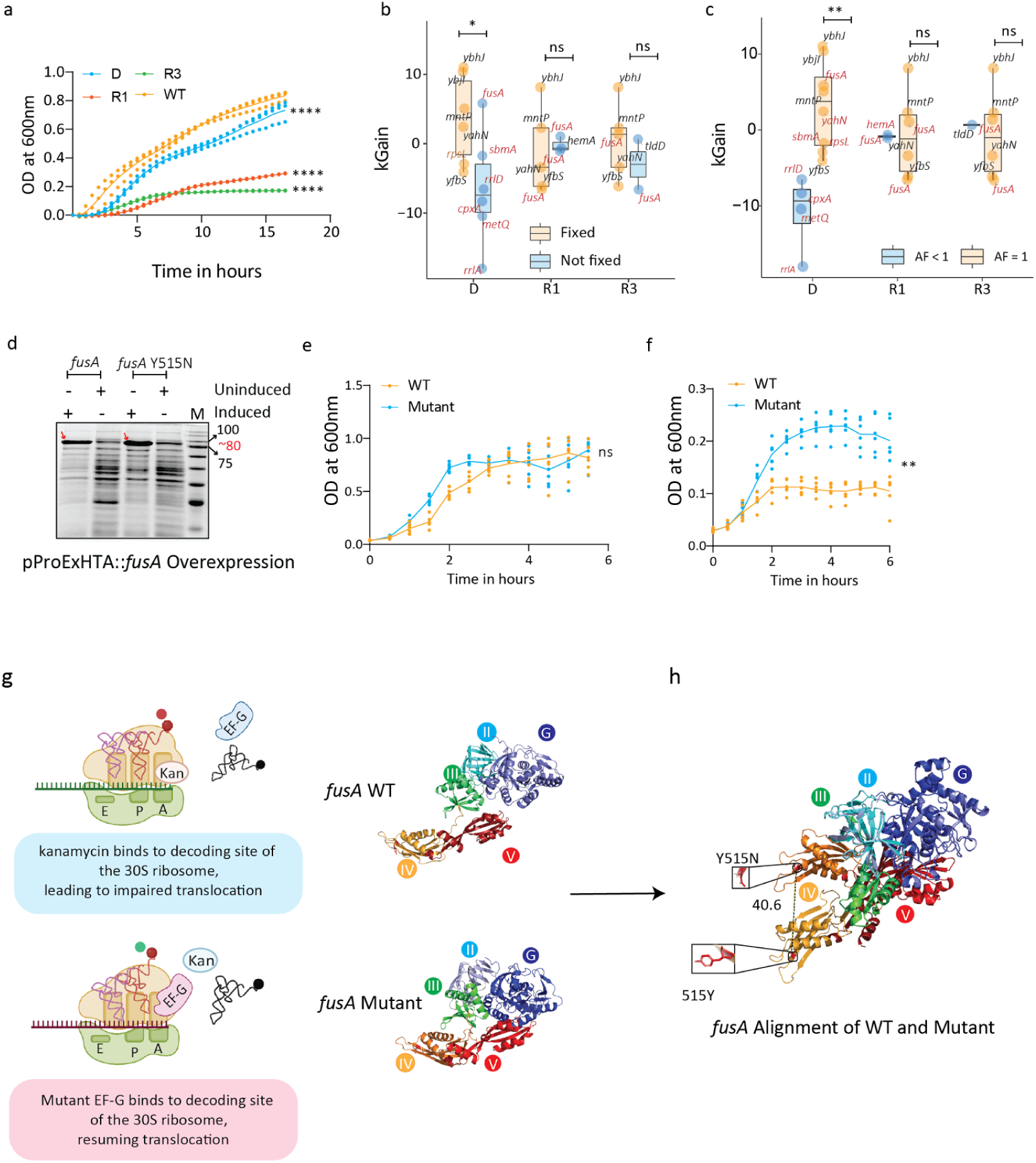
Impact of kanamycin and *fusA* mutations on ribosomal translocation. **(a)** Growth curve for passage 5 of the D, R1, R3 populations, and the WT founder strain in LB medium without antibiotics. The OD^600^ was measured over a 16-hour period to assess bacterial growth across all populations. **(b)** kGain scores for populations with fixed (orange) and not fixed (blue) mutations. Boxplots show the distribution of kGain scores for each population (D, R1, R3), comparing mutations that were successfully carried (*P*-value = 6.17e-01 and effect size = -1.05e+00 for R1, *P*-value = 1.87e-02 and effect size = 1.99e+00 for D, *P*-value = 2.58e-01 and effect size = 9.51e-01 for R3). **(c)** kGain scores for mutations in genes across populations D, R1, and R3 are shown . Boxplots depict the kGain scores for mutations in genes, comparing the alternate allele frequency (AF) of 1 (orange) versus AF less than 1 (blue) in each population (*P*-value = 4.84e-01 and effect size = -7.85e-02 for R1, *P*-value = 1.75e-03 and effect size = 2.53e+00 for D, *P*-value = nan and effect size = -4.00e-01 for R3). **(d)** SDS-PAGE analysis of induced and uninduced cells carrying pProExHTA::*fusA* and pProExHTA::*fusA* Y515N. BL21 cells transformed with pProExHTA::*fusA* and pProExHTA::*fusA* Y515N in log phase were separated into two tubes and one of each was induced with 1.0 mM IPTG at 37°C overnight. 20 µl of cells were mixed with SDS loading dye and boiled prior to loading in the gel. Red arrows indicate the expression of *fusA* or *fusA* Y515N in lanes loaded with induced cells. *M* denotes Marker in kDa. Growth curve for BL21 cells expressing *fusA* or *fusA* Y515N in the **(e)** absence or **(f)** presence of 0.008 mg/ml Kanamycin. The cells were induced with 0.4 mM IPTG and growth was followed and OD_600_ was measured for 8 hrs. **(g)** Schematic illustrating the proposed mechanism of ribosomal translocation in the presence of kanamycin (Kan). In the WT scenario, kanamycin binds to the decoding site of the 30S ribosome, leading to impaired translocation. In the *fusA* mutant, a mutant form of EF-G binds to the decoding site of the 30S ribosome, resuming translocation despite the presence of kanamycin. **(h)** Structural alignment of *fusA* WT and *fusA* mutant protein structures. The *fusA* WT structure is shown with regions I-V labelled and color-coded. The *fusA* mutant structure is aligned with the WT, highlighting the Y515N mutation, which is critical for resuming translocation in the presence of kanamycin.

Whole-genome sequencing of all clones enabled us to score all single-nucleotide variants (SNVs) by their kGain values using our established pipeline. In the D population, “fixed” mutations (persisting across ≥4 passages) bore significantly higher kGain than transient variants (**Fig. 6b; Supplementary Fig. S5c**), and mutations fixed at frequency = 1.0 exhibited elevated kGain relative to those that failed to fix (**Fig. 6c**; **Supplementary Fig. S5b**). Total SNV counts were similar across populations (D = 36; R1 = 29; R3 = 26), but D showed a pronounced skew toward high-frequency and fixed alleles, consistent with stronger selection.

To validate our hypothesis, we examined *fusA*, encoding elongation factor G (EF-G), a protein of five domains essential for ribosomal translocation. In population D, a Y515N substitution in domain IV emerged at passage 4 and fixed by passage 5 (kGain = 5.81). In contrast, R1 and R3 accumulated multiple *fusA* mutations in domains I and V, including I158N (kGain = –6.57), I654N (–1.06), T674A (1.05) in R1, and A76T (1.35), G575D (–6.60) in R3. To evaluate the functional consequences of the Y515N substitution in EF-G, we subcloned the wild-type and Y515N *fusA* alleles into the pProExHTA vector and transformed them into *E. coli* BL21(DE3). Expression of both wild-type and Y515N EF-G variants was assessed by SDS–PAGE under uninduced (0 mM IPTG) and induced (1 mM IPTG) conditions; the ≈80 kDa bands corresponding to each EF-G protein were observed only upon IPTG induction (**Fig. 6d**). In the absence of kanamycin, without IPTG induction,both strains exhibited identical growth kinetics, demonstrating no basal fitness defect. Upon induction and exposure to 0.008 mg·mL^-^ kanamycin, cells expressing *FusA* Y515N sustained robust growth over an 8-hour time course (**Fig. 6e**), whereas wild-type *FusA*, expressing cells, failed to proliferate. These results indicate that the Y515N mutation impairs kanamycin’s interaction with the ribosome, thereby conferring antibiotic resistance (**Fig. 6f**). Domain IV mutations have been implicated in modulating EF-G–ribosome interactions without abolishing translocation. Although EF-G is vital for translation, several studies have reported non-lethal *fusA* mutations in mediating aminoglycoside resistance (Holley et al. 2022; Mogre, Veetil, and Seshasayee 2017; Rodriguez de Evgrafov et al. 2020; Chulluncuy et al. 2016). While the precise mechanism remains unclear, we speculate that the Y515N substitution of *fusA* in domain IV affects its binding to the 30S ribosome (**Fig. 6g,h and Supplementary Fig. S5d**), preserving EF-G’s ability to facilitate translocation. Our overall findings suggest that k-mers are selectively favored even under selective pressure. In population D, robust growth was maintained, suggesting that Y515N could confer resistance without compromising fitness associated with higher kGain scores. In contrast, *fusA* variants R1 and R3 provided only partial resistance, correlating with lower kGain scores and diminished growth.

Together, these data demonstrate that k-mer based selection biases endure under antibiotic stress and can predict adaptive trajectories. The Y515N *fusA* mutation, characterized by a high kGain score, confers kanamycin resistance without compromising fitness, whereas lower kGain *fusA* variants in R1 and R3 afford only partial resistance at the cost of reduced growth. This work underscores the utility of kGain as a genome-wide metric for forecasting functionally important adaptive mutations.

### Experimental evolution in *S. cerevisiae* reveals consistent kGain trends

Following the assessment of a prokaryotic system, we inquired if eukaryotes would conform with the elevation of the kGain score levels along the evolutionary time-course. To this end, we performed meta-analysis of genomic sequencing dataset from a *S. cerevisiae* LTEE experiment ^33^. The study chose 30 focal populations (12 diploid populations, 12 MATa populations, and 6 MATα populations) across three environments, cultivation in yeast extract peptone dextrose (YPD) at 30°C, in synthetic complete medium (SC) at 30°C, and in synthetic complete medium (SC) at 37°C. We computed kGain scores for 133,538 substitutions from across the 90 populations under study. Notably, authors performed genomic sequencing for samples from only six representative timepoints. We observed a gradual increase in both the median fitness and median kGain scores over successive generations (**Fig. 7a**). To quantify this trend, we employed least squares regression and observed a strong correlation (Pearson’s *rho* = 0.9635) between median fitness and median kGain **(Supplementary Fig. S6a)**. Despite the absence of a rise in mutation numbers akin to those observed in the *E. coli* LTEE study, populations did not display hypermutator phenotypes, indicating a steady accumulation of beneficial mutations even in the absence of mutator phenotypes. We next examined the essential vs. non-essential genes dynamic and observed that essential genes consistently exhibited higher kGain scores compared to non-essential ones. This pattern implies a stronger selective pressure on essential genes to preserve beneficial or neutral variants. Notably, the trend remained significant across all generations, underscoring the greater evolutionary constraint these genes face in maintaining functional integrity (**Fig. 7b**). We closely examined the fixed mutations in the populations. The authors labelled a mutation as “fixed” at a particular time point if it met the criteria of having coverage of at least 5X and being present at a frequency of greater than or equal to 40% (for diploids) or 90% (for haploids). Notably, fixed mutations were assigned higher kGain scores, as compared to ones not fixed (**Fig. 7c**).

**Figure 7.**
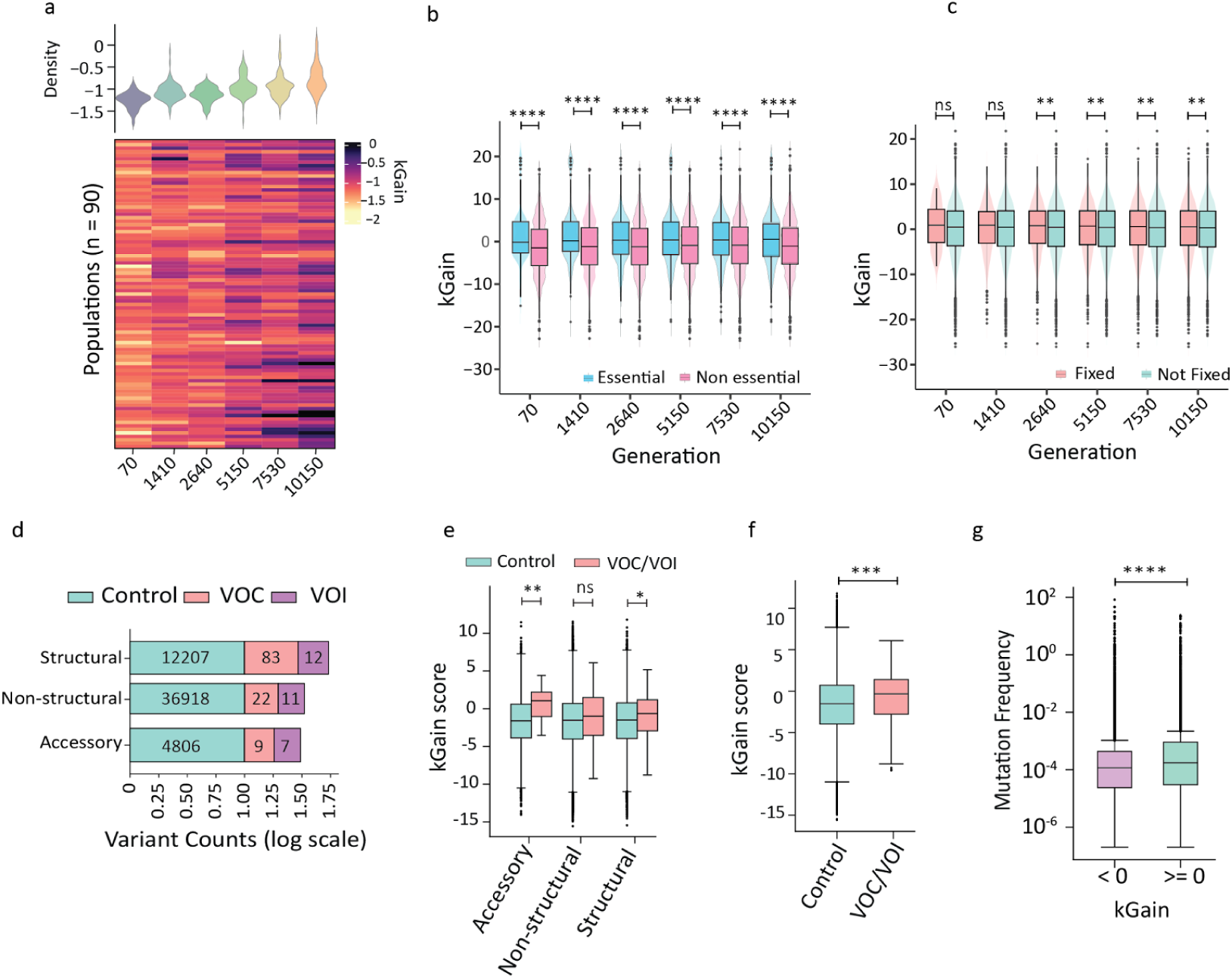
Evolution of kGain scores across generations and categories in Yeast and Covid. **(a)** Heatmap showing the distribution of median kGain scores across 90 populations at different generations, with marginal violin plots depicting the density of kGain scores across sampled generations in yeast. **(b)** Violin and box plots comparing kGain scores between essential and non-essential genes across unique mutations in yeast (*P*-value = 3.72e-16 and effect size = 3.06e-01 for generation 70, *P*-value = 1.69e-32 and effect size = 3.14e-01 for generation 1410, *P*-value = 1.39e-24 and effect size = 3.65e-01 for generation 2640, *P*-value = 1.52e-28 and effect size = 2.97e-01 for generation 5150, *P*-value = 6.82e-23 and effect size = 2.89e-01 for generation 7530, *P*-value = 1.04e-27 and effect size = 2.97e-01 for generation 10150). **(c)** Violin and box plots comparing kGain scores for fixed and not fixed mutations across unique mutations in Yeast (*P*-value = 1.73e-01 and effect size = 1.09e-01 for generation 70, *P*-value = 5.32e-02 and effect size = 1.19e-01 for generation 1410, *P*-value = 4.71e-03 and effect size = 9.49e-02 for generation 2640, *P*-value = 3.25e-03 and effect size = 8.53e-02 for generation 5150, *P*-value = 2.72e-03 and effect size = 6.00e-02 for generation 7530, *P*-value = 1.03e-03 and effect size = 6.08e-02 for generation 10150). **(d)** The variant counts are depicted using a logarithmic scale for control, VOC, and VOI categories within the structural, non-structural, and accessory gene groups in SARS-CoV-2. **(e-f)** All variants are classified into accessory, structural, and non-structural based on gene and variant classification (control and VOC/VOI) and then visualised with kGain using box plots in SARS-CoV-2. **(g)** A kGain of 0 is taken as a cutoff, and based on that, variants => 0 and variants <0 are visualised with the log of mutation frequency using a box plot in SARS-CoV-2.

### Pathogenic *SARS-CoV-2* variants are assigned elevated kGain scores

The recent outbreak of SARS-CoV-2 arguably offers the most extensive data on natural evolution, supported by comprehensive large-scale genomic profiling initiatives. To comprehensively analyze the mutational landscape of the SARS-CoV-2 virus, we implemented a robust pipeline for mutation analysis, as illustrated in **Supplemental Fig. S6b.** We took around 0.5 million high-quality genome sequences from GISAID (last accessed on: 29-08-2022) and made SNP calls, yielding ∼54,000 unique SNPs. Within our analysis, we delineated gene length and unique mutation counts **(Supplemental Fig. S6.c,d and table S1)**. We employed two widely recognized approaches to classify SARS-CoV-2 variants: functional categorization of associated genes and their pathogenic characteristics with implications for human health. Functionally, genes are divided into structural, non-structural, and accessory classes. Structural genes, such as Spike (S), Envelope (E), Membrane (M), and Nucleocapsid (N), are essential for the structural integrity and infectivity of the virus. These genes facilitate viral entry into host cells by binding to the ACE2 receptor and contribute to immune evasion and viral persistence. Non-structural genes, exemplified by ORF1ab, are crucial for viral replication and transcription which are vital for its lifecycle and interaction with the host immune system. Accessory genes, including ORF3a, ORF6, ORF7a, ORF7b, ORF8, and ORF10, enhance the ability of the virus to evade immune responses and modulate host cell functions such as antagonizing the host’s interferon response (**Fig. 7d**). Based on pathogenic roles and public health implications, the World Health Organization (WHO) has classified SARS-CoV-2 variants as Variants of Interest (VOI) or Variants of Concern (VOC) **(table S2)**. VOIs have mutations affecting virus behavior, including transmissibility, severity, detection, or treatment. VOCs meet VOI criteria and cause severe disease, significantly reducing vaccine effectiveness and impact healthcare systems. Examples of lineages associated with VOIs include lineages like Lambda and Mu, while those associated with VOCs include Alpha, Beta, Gamma, Delta, and Omicron **(Supplemental Fig. S6e)**. More details on COVID-19 data processing and filtering can be found in the Materials and Methods section.

We observed a significant contrast between the control variants and the VOC/VOI variants, with the latter exhibiting substantially higher kGain scores (**Fig. 7f**). Interestingly, gene groups revealed that kGain scores for VOC/VOI variants were higher than those for control variants, particularly within accessory and structural genes (**Fig. 7e**). We set the kGain cutoff at 0 and divided the variants into two categories: <0 and ≥0 (**Fig. 7g**). Variants in the latter group were found to be present in considerably higher numbers of COVID-19 genome sequences, suggesting that our score could indicate the type of selection pressure (positive or negative) experienced by a mutation.

Zhang et al. (2022) ^34^ reported mutations that contribute to the pathogenicity and fitness of the virus. We examined the kGain profile of the reported VOC/VOI mutations and found that ORF3a:S26L, ORF3a:T223I, and ORF3a:S171L had higher kGain scores, whereas ORF3a:Q57H and ORF3a:S253P had lower kGain scores. These variants highlight changes in accessory proteins, which may exhibit a heightened propensity for adaptation through immune evasion.

Another study by Obermeyer et al. (2022) ^35^ established that the SARS-CoV-2 pandemic has been dominated by several genetic changes of functional and epidemiological importance, including the spike (S) D614G mutation. This mutation is associated with higher SARS-CoV-2 loads and has contributed to the increased infectivity and fitness of the virus **(Supplementary Note 8)**

### Deep Mutational scans (DMS) uncover kGain signatures in key genes

Having demonstrated kGain’s utility in capturing evolutionary constraints in both the *E. coli* LTEE and our in-house antibiotic adaptation experiment, we next applied it to deep mutational scanning (DMS) data. DMS systematically evaluates all possible substitutions in essential genes, providing a comprehensive landscape of mutational effects. By examining kGain scores in this context, we aimed to determine whether the adaptive signals observed in long-term evolution also arise under the focused conditions of DMS.

We analysed three essential *E. coli* genes *murA*, *fabZ*, and *lpxC*, each vital for cell envelope biosynthesis ^36^. Specifically, *murA* catalyzes the first step in peptidoglycan precursor formation, *fabZ* acts as a dehydratase in fatty acid synthesis, and *lpxC* produces lipid A, a core component of the lipopolysaccharide (LPS) layer. Among these, *murA* appears significantly mutation-sensitive, potentially permitting more beneficial shifts than *fabZ* or *lpxC*. A heatmap (**Fig. 8a,b**) comparing kGain with tolerance scores showed alternating gradients, where regions of high tolerance often correlated with higher kGain. Similar patterns emerged for *lpxC* and *fabZ* (**Supplemental Fig. S7a,b**). By applying a tolerance-score cutoff of 0.8 (**Fig. 8c,e**), we observed that “tolerant” mutants generally had higher kGain than “non-tolerant” ones (cutoff = 1, **Supplemental Fig. S7c,e**), implying that some variants could impact beneficial effects.

**Figure 8.**
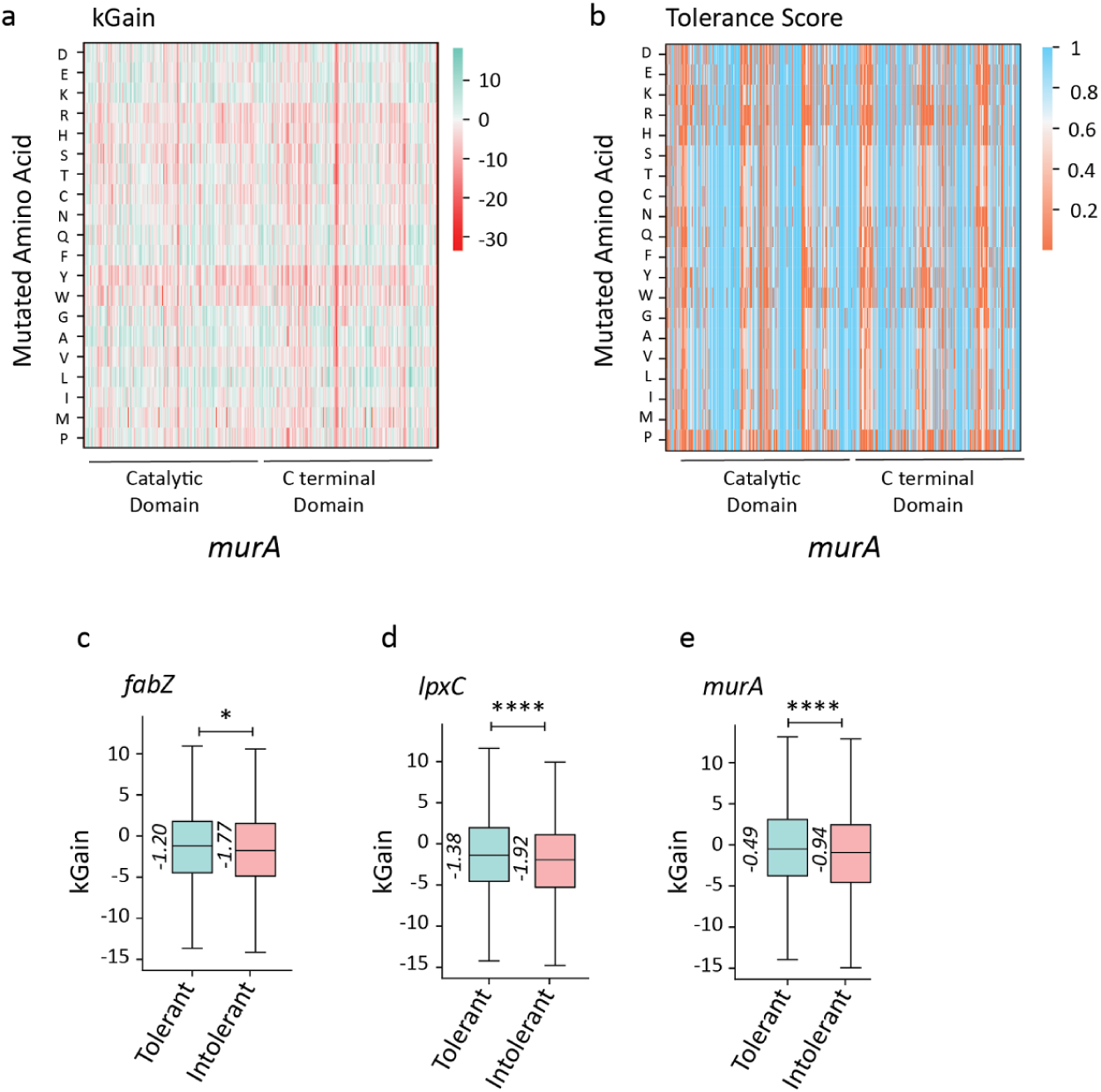
kGain scores dynamics captured in DMS studies. **(a)** Heatmap showing the median kGain scores for mutations across different amino acids in *murA* gene. The color scale indicates the range of kGain scores, with red representing negative values and green indicating positive values. **(b)** Heatmap representing the median tolerance scores for each mutated amino acid, with the color scale from blue to red representing the range of tolerance values from 0 to 1. The domains are categorized as catalytic and C-terminal, with distinct coloring to highlight regions of higher or lower tolerance. **(c-e)** Boxplot showing kGain scores for *fabZ, lpxC*, and *murA* mutations, comparing tolerant versus intolerant mutations. Tolerant mutations (green) are associated with higher kGain scores, while intolerant mutations (red) show significantly lower values (*P*-values are 2.08e-2 for *fabZ*, 7.11e-9 for *lpxC*, and 5.46e-8 for *murA*). [**Note**: The p-value cutoff for all the plots is 0.05. *, **, ***, and **** refers to p-values <0.05, <0.01, <0.001, and <0.0001, respectively.]

To explore the functional impact of these mutations, we examined relative solvent accessibility (RSA) in conjunction with kGain. RSA indicates whether a residue is surface-exposed or buried within the protein structure. We found that residues with RSA > 1 (i.e., surface-exposed) frequently aligned with higher kGain (notably in *murA* and *lpxC*) **(Supplemental Fig. S7f**), suggesting that exposed sites may be more prone to accumulate beneficial mutations.

## Discussion

This study introduces kGain as a compact and interpretable metric that captures selection-linked shifts in oligonucleotide frequency during evolution. Traditional models of selection often rely on changes in protein structure, interspecies conservation, or allele frequency trajectories. Here, we propose that within-genome k-mer dynamics offer an orthogonal and quantifiable dimension of evolutionary constraint and selection.

Across diverse evolutionary systems, including laboratory-evolved *E. coli* and *S. cerevisiae* populations, SARS-CoV-2 genome surveillance, and in-house antibiotic adaptation experiments, we observed a consistent trend: variants with higher kGain scores tend to persist and fix over time. These findings suggest that high-kGain contexts, reflecting enriched sequence motifs, are more likely to harbor beneficial or tolerated mutations. This is particularly evident in essential genes, where elevated kGain scores imply selective pressure not only on the coding potential but also on local sequence architecture that may influence regulatory robustness or mutational resilience.

Our analysis reveals that fixed variants, essential-gene mutations, and substitutions in parallel genes all show higher kGain scores than their respective controls. Parallel genes, recurrently mutated across independent LTEE populations, are especially informative. We find that these genes accumulate mutations more rapidly and in higher-kGain contexts, particularly in mutator backgrounds. Bootstrapped trajectories further reveal a generation-wise increase in both allele frequency and kGain in parallel loci, indicating that recurrent adaptive evolution may be facilitated by favorable motif architectures. These results suggest that mutators, despite their elevated mutation load, are not phenotypically compromised because they sample and fix beneficial mutations in high-kGain sequence contexts. This provides a mechanistic rationale for the fitness parity observed between mutator and non-mutator lineages over 60,000 generations.

In the context of environmental stress, our single-colony bottleneck experiment under sublethal kanamycin selection showed that fixed mutations in the adaptive lineage (Population D) were significantly enriched for high-kGain scores. The *fusA* Y515N mutation, which reached fixation in Population D, conferred kanamycin resistance without growth cost when overexpressed in a wild-type background. In contrast, *fusA* mutations with lower or negative kGain in other populations failed to confer resistance or sustained growth. These results provide direct functional validation that high-kGain contexts are predictive of adaptive potential.

Complementing our experimental validation, we demonstrate that kGain is computationally predictable. A transformer-inspired attention-based neural network, trained on short nucleotide windows flanking SNPs, achieved a Pearson correlation of r = 0.82. The model learned biologically meaningful patterns from both the local sequence context and the mutant allele, suggesting that k-mer-driven selection pressures can be approximated without exhaustive full-genome frequency computation. Importantly, this also opens the door for applying kGain inference across species using learned representations.

Our DMS analysis further supports these insights. While the correlation between kGain and tolerance scores is modest, we observe that residues with high relative solvent accessibility (RSA), especially in *murA*, tend to show elevated kGain. This suggests that surface-exposed residues may be more permissive to beneficial substitutions in enriched k-mer contexts. Although kGain alone is not a substitute for functional assays, it captures a meaningful dimension of mutational tolerance that may complement structural and biochemical predictors.

We also observed significant kGain enrichment in natural viral evolution. SARS-CoV-2 Variants of Concern (VOCs) and Variants of Interest (VOIs) show elevated kGain scores, especially in structural and accessory genes. This suggests that even in fast-evolving RNA viruses, motif-level constraints may influence the success and spread of adaptive mutations. These findings align with prior work demonstrating that motif-level conservation is preserved even in genomes under rapid drift.

While our study focuses on point mutations, we acknowledge that other forces, horizontal gene transfer, indels, transposons, also shape genome evolution. However, the controlled context of LTEE and the reproducibility of kGain enrichment across independently evolved systems argue for a consistent selection-based mechanism at the k-mer level.

In conclusion, our findings support a model in which selection not only targets genes or phenotypes but also operates on the sequence context itself. The consistent enrichment of high-kGain mutations in adaptive lineages, essential loci, and parallel genes, combined with predictive accuracy from machine learning, suggests that k-mer architecture plays a previously underappreciated role in shaping evolutionary trajectories. As genome-wide data continues to accumulate, incorporating motif-level selection into evolutionary models could provide new insights into how mutation, sequence context, and fitness co-evolve.

## Methods

### Mathematical formulation of FCGR

Let ***S*** be a DNA nucleotide string of length **N**, where ***S***[*i*] represents the *i*^th^ symbol (1≤*i*≤N) corresponding to a DNA nucleotide base (A, C, G, T). The notation ***S***[..*i*] denotes the DNA sequence prefix ending at position *i* (***S***[..*i*]=***S***[1..*i*]). The CGR iterative algorithm operates on an ℝ^2^ space, with each vertex corresponding to a DNA base (A, C, G, T). For a given DNA sequence ***S*** of length **N**, CGR maps each ***S***[..*i*] prefix to the point *x^i^*∈ℝ^2^ using an iterative procedure.

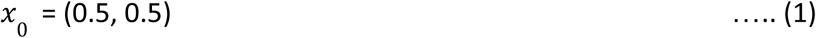

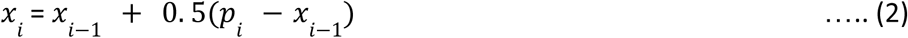

Where,

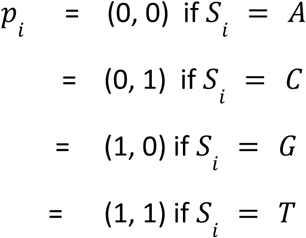

In the original formulation ^37^, the initial point (*x_0_*) was taken as the center of the square (0.5, 0.5). Alternatively, this point could be chosen randomly within the square. The user can also customize the vertex for *p_i_* corresponding to a given *S_i_*. Here, *S_i_* represents the randomly chosen vertex for the *i^th^* step of the walker.

### FCGR heatmap generation

We used the kaos Python package ^19^ to produce the FCGR encoding matrix. The *k*-mers frequencies, returned by kaos, were scaled by dividing each *k*-mer frequency by the total *k*-mer count in the DNA sequence. The 1024 x 1024 pixel FCGR heat maps show the negative log of these normalized frequency values.

### Computing the kGain score

kGain scores associated with single nucleotide variation (SNVs) are computed solely by using the reference genomes of the respective organisms – *E. coli B str.* REL606 for the Goods dataset, *S. cerevisiae* for budding yeast LTEE and *Severe acute respiratory syndrome coronavirus 2* isolate Wuhan-Hu-1 for *SARS-CoV-2*. Further details pertaining to the individual datasets can be found in the following sub-sections.

Below are the steps for kGain score computation.

**A. Sequences with the SNV and flanks:** For every variant, we generate two sequences of length 19 (considering *k*-mer length is 10), one with the variant allele at the middle (10^th^ position), and the other with the reference allele at the same location. Left and right flanks are sourced from the associated reference genome, depending on the organism.
**B. *k*-mer generation using rolling windows:** A total *k* (*k* = 10) windows are generated for each variant, such that each *window* contains the reference/alternate allele i.e., the position of the alternation. Rolling window based *k-*mer generation is performed in pair once for the reference sequence and once for the sequence carrying the variant of interest, giving rise to sets of k-mers per variant.
**C. kGain computation:** For each *k-*mer, its occurrence across the reference genome is tracked, which finally contributes to the kGain computation.

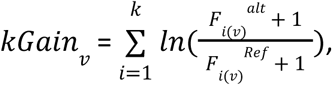

Where, *kGain_v_*, score is computed for each variant *v* by adding the natural log of the fold change between the genomic frequencies of the *k*-mer containing the alternate allele (*F*_i(v)_*^alt^*) and the reference allele (*F*_i(v)_*^ref^*) within each i^th^ window, across all windows. In this equation *k* is the total number of *k-*mers generated using the rolling window method, both for the reference and alternate sequences. Since the computation is on a logarithmic scale, both division by zero and taking the logarithm of zero pose issues. To address this, a pseudo count of 1 is added to both the numerator and denominator. This ensures numerical stability by preventing division by zero and avoiding undefined logarithmic values when the alternate k-mer frequency is 0.

### LLR Score

We utilize the ESM1b model to compute the Likelihood ratio (LLR) exclusively for missense mutations across all datasets described below. The ESM1b model is being harnessed from the GitHub repository (https://github.com/ntranoslab/esm-variants).

### Evo Score

The Evo model, trained on 2.7 million prokaryotic and phage genomes, allows zero-shot prediction of how small nucleotide sequence changes affect overall organismal fitness. For each mutation, we extracted a 101-base sequence centered on the mutation position, including 50 bases upstream and 50 bases downstream, with the mutated allele at the center. The Evo-1-131k-base model was then used to generate a log likelihood score for each 101-base sequence (https://github.com/evo-design/evo).

### Overview of dataset

To explore the utility of kGain score, we performed meta-analyses of genomic sequencing data from laboratory evolution experiments involving *E. coli* and S. *cerevisiae* (budding yeast). Further we used *Severe acute respiratory syndrome coronavirus 2* (*SARS-CoV-2*) data from the GISAID repository. Apart from these, datasets from two other studies were used for various analyses.

### *E. coli* LTEE data (Goods dataset)

The *E. coli* LTEE is an ongoing study in experimental evolution that began in 1988 by Richard Lenski at the University of California, Irvine ^38^. It observes the evolutionary alterations across 12 genetically identical populations that have been consistently maintained in a controlled environment ^39^ ^26^. We collected bacterial fitness measurements and sequencing data from two independent studies by Weiser et al. and Goods et al. Weiser and colleagues performed competitive fitness assays for up to 50,000 generations, totalling 928 screened samples across 12 populations. 41,962 variants identified by Goods and colleagues ^30^ were filtered to include only single nucleotide substitutions, resulting in 36,923 variants. Both these datasets are accessible at https://github.com/benjaminhgood/LTEE-metagenomic.

### *S. cerevisiae* LTEE data (Johnson dataset)

The S. *cerevisiae* LTEE experiment comprised 205 (124 haploid and 81 diploid) populations, propagating up to ∼10,000 generations in three different environments ^33^. For each of the three environments, the authors selected 30 focal populations (12 diploid, 12 *MATa*, and 6 *MAT*α) and sequenced them at six-time points, giving rise to 90 whole genome sequences. Upon request, we received the unfiltered *.vcf* file (144,708 variants), from which we filtered the single base substitutions, resulting in 133,538 variants. We obtained the fitness data from the same publication ^33^.

### SARS-CoV-2 dataset

About 0.5 million (N=477, 667) high-quality *SARS-CoV-2* genomic sequences were downloaded from the Global Initiative on Sharing All Influenza Data (GISAID) (GISAID Initiative (epicov.org)) EpiCoV database (last accessed on 29-08-2022) based on stringent criteria - human as host, completed genomes, excluding low coverage data, availability of patient clinical status and collection date ranging between 1st Jan 2010 and 29th Aug 2022. Genome-wide variant identification and open reading frame (ORF) prediction were executed using reference genome (Accession ID: NC_045512.2), employing an intricate multi-step analytical approach that integrated the StrainFlow and CoV-Seq pipelines as described by the authors. From a total of 68,364 unique mutations, 56,971 were identified as SNPs. Among these, 54,085 SNPs were considered for further analysis. At the same time, 2886 variants were excluded due to the presence of common variants spanning multiple genes, flank lengths less than ten bp and lacking protein variant identifiers (HGVS_P). The refined variant dataset was provided as input for the computation of the kGain score, as detailed above.

### *SARS-CoV-2* dataset preprocessing

The preprocessing of the dataset contains the following steps :

1. **Calculating the mutation frequency and mutation count**: The mutation frequency for 54,075 SNPs is determined by dividing the mutation counts by the total sum of mutation counts within a gene. These mutation counts for each variant were obtained from the Python Outbreak.info API package. Notably, 10 SNPs were excluded from the analysis as they were not accessible through the API (Welcome to the Python Outbreak.info package docs! — Python Outbreak.info API 0.1 documentation (outbreak-info.github.io)).
2. **Integration of Lineage and Variant Classification in *SARS-CoV-2***: Data extracted from the Python Outbreak.info API comprised 334 variants, each associated with lineage information including alpha, beta, gamma, delta, omicron, b.1.2, lambda, and mu. Integrating this lineage data with unique SNPs allowed us to identify 144 variants. Following this identification, these 144 variants were categorized into variants of concern (VOC) and variants of interest (VOI), in accordance with the SARS-CoV-2 variant classification guidelines established by the WHO (COVID-19 variants | WHO COVID-19 dashboard and SARS-CoV-2 Variant Classifications and Definitions (cdc.gov)). The remaining variants were classified as controls within the analysis.

### Bacterial adaptation experiment

#### Bacterial Strains

*Escherichia coli (E. coli)* strain MG1655 was used as the wild-type (WT) founder for the adaptation experiment. All strains employed were isogenic with *E. coli* MG1655.

For clarity, the following definitions apply: **Variants/Mutations:** The list of mutations that distinguish a mutant from its isogenic parent strain. **Lineage:** A temporal series of bacteria connected by a continuous line of descent from ancestor to descendant. **Clone:** A population of bacteria (typically a colony on agar) derived from a single founder cell. **Founder:** WT bacterial strain used as a starting point for the experiment.

#### Cloning and Site Directed Mutagenesis

*fusA* gene was cloned using standard cloning techniques. Briefly, *fusA* gene was PCR amplified using the primer pair: Forward Primer - 5’ TATA<u>GGATCCA</u>ATGGCTCGTACAAC 3’ and Reverse Primer - 5’ CTTTT<u>CTCGAG</u>TTATTTACCACGG 3’. PCR product and pProExHTA vector backbone were digested using BamHI and XhoI to generate linear DNA fragments with sticky ends, ligated, and transformed in DH5⍺ cells. Recombinant colonies were screened and sequencing was done to confirm that *fusA* was free of any spontaneous mutations.

pProExHTA::*fusA* was used to generate pProExHTA::*fusA Y515N* using Phusion Site Directed Mutagenesis Kit (ThermoScientific #F541) and primers: Forward Primer- -- 5’ GTCGTGGTCAG**<u>A</u>**ATGGTCATGTT 3’ and Reverse Primer-5’ AACATGACCATTCTGACCACGAC 3’.

#### Culture conditions

Two types of media were used for the adaptive experiments: Luria–Bertani (LB) broth/Luria Agar (LA). For 100 mL of LB, the medium was composed of 1 g tryptone, 1 g NaCl, 0.5 g yeast extract. The LA consists of LB medium with 1.5g agar. The pH was adjusted to ∼7.0 prior to autoclaving at 121 °C for 15–20 minutes. M9 Minimal Medium (M9A): M9 minimal medium was prepared using a 5× M9 salts solution containing 32 g Na_2_HPO_4_, 7.5 g KH_2_PO_4_, 1.25 g NaCl, and 2.5 g NH_4_Cl dissolved in 500 mL of distilled water. For 500 mL of M9A, the salts were combined with additional components, including 1 M MgSO_4_, 1 M CaCl_2_, 20% glucose, agar, and 250 µL of trace elements solution (comprising FeCl_3_, ZnSO_4_, CuCl_2_, MnSO_4_,and CoCl_2_).

#### Bacterial adaptation under selective pressure: Experimental setup

Four independent lineages were established from the WT. Population D was propagated on LA plates, whereas Populations R1, R2, and R3 were maintained on M9A ^40,41^. A serial passage strategy was implemented, whereby at each passage a single colony was selected and transferred to fresh plates containing an incrementally increased sublethal concentration of kanamycin ^42,43^. Colony for each passage was based on selection and was tailored to the specific growth environment: in LA, colonies were chosen based on growth curve analysis ^44^—since rapid proliferation in a nutrient-rich medium renders growth kinetics a robust indicator of fitness—while in M9A, the INT/PMS assay was used to assess metabolic activity ^45,46^, which more accurately reflects cellular fitness under nutrient-limited conditions.

Following selection, the selected colony underwent minimum inhibitory concentration (MIC) determination ^47^, and the sublethal concentration thus established was used for the subsequent propagation. Initial MICs for kanamycin were 0.006 mg/ml in LA and 0.01 mg/ml in M9A, reflecting distinct metabolic conditions. LA’s nutrient-rich environment promotes rapid growth, potentially increasing antibiotic susceptibility, whereas M9A requires de novo synthesis of essential metabolites, resulting in slower growth and a slightly higher MIC ^48,49^). For Population D, kanamycin concentrations escalated from 0.006 mg/ml to 0.008, 0.01, 0.04, and 0.08 mg/ml across D1–D5. In M9A, Population R1 encountered 0.01, 0.05, 0.06, and 0.2 mg/ml, while Population R3 followed a similar progression, culminating at 0.1 mg/ml by passage 5 (**Supplementary Note 6**).

#### Colony Selection via Divergent Fitness Assays

Colony selection criteria were tailored to the growth environment; for population D (LA), colony selection was based on growth kinetics. A 1% inoculum from a 3-hour culture (OD_600_ ∼0.8–1.0) was added to a 96-well plate containing fresh LB. Growth was monitored for 16 hours with OD_600_ measured at ∼ 30-minute intervals. The area under the curve (AUC) was calculated, and colonies exhibiting the highest AUC were selected for the next passage.

Populations R1 and R3 (M9A): Due to the nutrient-limited conditions, metabolic activity was assessed using an INT (1 mM) plus PMS (2.5 mM) redox-based colorimetric assay. Cultures grown to an OD_600_ of 0.8–1.0, 10 µL aliquots were applied onto Whatman® grade 3 filter papers. Following the addition of 5 µL of the INT/PMS solution, the development of a purple formazan precipitate was monitored. Samples were air-dried in the dark (1–4 hours), scanned, and the color intensity (measured in the green channel) quantified using ImageJ™. Colonies with the highest metabolic activity were advanced to the subsequent passage.

#### Antibiotics and Sublethal Concentration Selection

Kanamycin (purchased from SRL Labs, India) was dissolved in nuclease free water at a concentration of 50 mg/ml prior to use. Kanamycin (Kan) was added to liquid and solid media at the specified concentrations . To determine the appropriate sublethal antibiotic concentration for each passage, minimal inhibitory concentration (MIC) assays were performed in 96-well plates. Briefly, 1% of a pre-cultured bacterial suspension (OD_600_ between 0.8 and 1.0, following a 3–4-hour incubation) was inoculated into wells containing progressively increasing concentrations of kanamycin. Optical density (OD_600_) readings were recorded 18–20-hour period at 37 °C.

#### Whole-Genome Sequencing

Whole genome sequencing (WGS) was outsourced where genomic DNA was extracted from colonies using the DNeasy PowerSoil Pro Kit (Qiagen) according to the manufacturer’s protocol. Library preparation was followed by paired-end sequencing (150 bp) on an Illumina NovaSeq 6000 platform. Raw reads were quality filtered with fastp v0.12.4, and aligned to the *E. coli* K-12 MG1655 reference genome (ASM584v2; GCF_000005845.2) using BWA-MEM^50^. Reads were converted to BAM format using samtools v1.6, sorted with sambamba v1.0.1, and duplicate reads were marked. Variant calling was performed with FreeBayes v0.9.21.7 after base quality recalibration with htslib v1.21. Variants were filtered using vcflib v1.0.10 and vt v2015.11.10, applying a minimum quality threshold of 20 and a minimum total read depth of 10. Quality control metrics were generated using MultiQC v1.19, and the final variant set was manually annotated using a GFF file (GCF_000005845.2_ASM584v2_genomic.gff) to retrieve the names of the genes.

#### Growth Comparison Assays

To assess growth differences among evolutionary passages and populations, standardized growth assays were performed in antibiotic-free LB medium. Overnight cultures (grown without antibiotic to allow recovery) were diluted 1% into fresh LB and grown to an OD_600_ of ∼0.8. Subsequently, 1% inocula were transferred into 96-well plates containing LB, and growth was monitored for 16 hours with OD_600_ readings every ∼ 30 minutes. This approach enabled a comparative evaluation of growth kinetics across different passages and populations.

For growth comparison assay between WT and Mutant *FusA*, plasmids (pProExHTA) carrying the *fusA* and *fusA* Y515N genes were transformed into E. *coli* BL21 (DE3) cells. A single colony of each was inoculated and grown overnight. 1 % of primary inoculum was used for 5 ml of secondary inoculation and grown for 3-4 hrs, till OD_600_ reached 0.6. Subsequently, a 96-well plate containing 0.4 mM IPTG and 0.008 mg/ml kanamycin was set up with culture OD_600_ of 0.05. Growth was followed for 8 hrs and OD_600_ was recorded and plotted every 30 minutes.

### Deep Mutation Scan Dataset

Deep Mutational Scanning (DMS) data ^36^ we computed kGain scores for all edits for *lpxC, fabZ*, and *murA* genes, which are essential for bacterial viability. kGain was computed as mentioned above. Unlike typical nucleotide mutations, DMS involves codon mutations. Therefore, we adjusted our methodology to ensure the mutated codon position was always present within the sliding window used for kGain calculations. This involved generating flanking sequences to maintain the full three-nucleotide codon within every window.

Consequently, both the reference and alternate flanks in our DMS analysis were standardized to a length of 17 nucleotides.

### Statistical analysis

Throughout the paper, we have used one-sided Mann-Whitney U tests for statistical significance analysis between two groups, wherever applicable, unless specially mentioned. The one-sided Mann-Whitney U test is a non-parametric test used to assess whether one data group tends to have larger values than the other, with the effect size calculated as described in **Supplementary Note 9**.

### Sequence Embedding Strategy for SNP Effect Prediction

Accurate prediction of mutational effects requires that a model recognize both the specific nucleotide change and its sequence neighbors. However, standard one-hot encoding (OHE) for SNPs often fails to capture the precise position or nature of the mutation, limiting both biological interpretability and predictive power. To overcome this limitation, we introduced a dual-encoded sequence representation. For each SNP, a sequence window (length: 2*kmer length - 1) centered at the mutation was extracted. At each position, nucleotides were encoded using standard one-hot vectors for A, C, G, and T. At the central (mutated) position, both the reference and alternate alleles were explicitly encoded, with the alternate nucleotide assigned a negative value in its channel. This approach highlights the mutational event within the embedding, while also encoding wild-type and absent states. By providing the model with information on both the sequence neighborhood and the explicit mutation, the dual-encoded representation enables more accurate and interpretable predictions of mutational effects, outperforming standard encodings in this context.

### Focal Loss Optimization for Accurate SNP Effect Prediction

In biology, a single nucleotide change can have different effects depending on its location in the genome. Some mutations drive adaptation through directional selection, some are eliminated by purifying selection, and many simply drift without strong effect. Because the impact of each SNP is so context-dependent, predicting mutation effects can be especially challenging for rare or complex cases. To address this, we combined our biologically informed SNP embedding with a focal regression loss (gamma = 2), which places greater emphasis on large prediction errors. This strategy encourages the model to focus on difficult and rare SNPs, not just those that are easy to predict. By integrating context-aware embeddings and focal loss optimization, this approach delivers more accurate and robust predictions across the full range of mutation outcomes in the population.

### Attention-Based Neural Network for kGain Prediction

We implemented a deep learning regression model inspired by the transformer architecture to predict quantitative mutation effects from DNA sequence context. Each input is a Dual-Encoded Sequence Representation (19X4), a one-hot matrix spanning 19 nucleotides centered on the SNP. The architecture begins with a dense layer that projects each nucleotide position into a continuous embedding space. Learnable positional embeddings are then added to retain information about nucleotide order within the sequence. The combined embeddings are then processed by two multi-head self-attention layers (each with four heads), which enable the model to integrate information from all positions in the sequence, capturing important patterns and relationships between nucleotides. Each attention layer is followed by dropout regularization, residual connections, and layer normalization to stabilize learning and prevent overfitting. After attention, the sequence representations are further refined by a position-wise feed-forward block (with ReLU activation and dropout), again followed by residual connections and normalization. A global average pooling layer aggregates the information across all sequence positions, producing a fixed-length vector. This is passed through a final dense layer to generate a single, continuous output value representing the predicted effect of the SNP. All hyperparameters, including sequence length, embedding dimension, attention heads, feed-forward dimension, number of layers, and dropout rate, were optimized for robust performance and reliable convergence on both training and validation data.

### Logistic Regression for Predictors of Directional Selection

We modeled whether a mutation experienced directional selection (1) versus all other regimes (0; purifying selection or drift) as a function of mutation type (is_AT_to_GC, 1 = A/T→G/C, 0 = other) and kGain score using logistic regression. Regression coefficients (excluding the intercept) were exponentiated to obtain odds ratios, representing the change in odds of a mutation falling under directional selection per unit increase in each predictor.

## Acknowledgements

We would like to express our gratitude to Dr. Michael Desi from the Department of Organismic and Evolutionary Biology at Harvard University, Cambridge, and Dr. Milo Johnson, Postdoctoral Fellow at UC Berkeley, for their invaluable assistance in understanding and acquiring the data. We thank Dr. Manjula Reddy and Suraj Kumar Meher from the Centre for Cell and Molecular Biology (CCMB), Hyderabad, for the bacterial strain *E. coli* MG1655.

## Authors contributions

Conceptualization: DS. Methodology: DS, BM, AH. Investigation: BM, AH, SP, NJ, DP, ST, AG. Visualization: BM, AH, SP. Supervision: DS, GA, VM. Writing – original draft: DS, BM, AH. Writing – review & editing: DS, BM, AH, SMP, SP, AG, GA, VM, SS.

## Data and materials availability

The kGain analysis code and datasets for both the *E. coli* LTEE and *S. cerevisiae* LTEE, as well as the bioinformatic pipelines (alignment, variant calling, and annotation) for the *E. coli* MG1655 experiment and DMS studies, are publicly accessible on GitHub https://github.com/cellsemantics/Oligo_Promotion. The raw reads for the *E. coli* antibiotic adaptation generated in this study has been deposited in the NCBI database under the BioProject accession number PRJNA1225006. The kGain analysis code along with the attention model and data for *SARS-CoV-2* are available on Github https://github.com/cellsemantics/Oligo_Promotion. GISAID sequence data for *SARS-CoV-2* is publicly available at GISAID Initiative (epicov.org). Source code for variant calling, ORF prediction, and genome annotation is available at rintukutum/strainflow_manuscript (github.com) and boxiangliu/covseq: CoV-Seq: COVID-19 Genomic Sequence Database and Visualization (github.com). Reference genomes for *SARS-CoV-2* are sourced from Severe acute respiratory syndrome coronavirus 2 isolate Wuhan-Hu-1, co - Nucleotide - NCBI (nih.gov). The mutation count and lineage data are available at Welcome to the Python Outbreak.info package docs! — Python Outbreak.info API 0.1 documentation (outbreak-info.github.io). WHO guidelines for *SARS-CoV-2* variant classification are available at COVID-19 variants | WHO COVID-19 dashboard and SARS-CoV-2 Variant Classifications and Definitions (cdc.gov).

