## Supplemental Material for "Intra-DNA k-mer Conservation Patterns Encode Evolutionary Selection of Variants"

[#,†]Equal contribution

**Supplementary Note 1. Choice of k in kGain computation**

To compute kGain, choosing the appropriate k-mer length is crucial. We analysed how the number of distinct k-mers varies with k-mer length and found k = 10 as a change point. This choice also aligns with earlier studies (Ferreira et al. 2022). Consequently, we used k = 10 for all subsequent analyses (**Supplemental Fig. S1e,f**).

**Supplementary Note 2. Classification of Mutational Trajectories by Selection Regime**

Mutational trajectories were classified into directional selection, purifying selection, or drift by modeling the rate of change in allele frequency over time. For each mutation observed in at least ten generations, we carried out the following steps:

1. Log-odds transformation

   Denote the allele frequency of a given mutation for a given population at generation $t$ by $p_t$. Since allele frequencies lie between 0 and 1, we use the logit transformation to map these probabilities to the real number line. This transformation is essential for applying linear regression, because the linear predictor can take any real value, while probabilities are bounded between [0,1]. By converting data to the log-odds scale, we ensure compatibility between the data and the linear model framework.

   $$y_t = logit(p_t) = \ln(\frac{p_t}{1-p_t})$$

   Linear regression of log-odds across generation

   We fit a linear ordinary least-squares regression model using scipy.stats.linregress to predict $y_t$ as a linear function of generation number ($t$), and we note the estimated slope ($\beta_1$) and the two-sided p-value.

   $\beta_1$ represents the per-generation change in the log-odds of the mutant allele.

2. Statistical classification
   - **Directional selection**: $\beta_1 > 0$ with two-sided $p < 0.05$

- **Purifying selection**: $\beta_1 < 0$ with two-sided $p < 0.05$
- **Drift/Neutral**: two-sided $p > 0.05$ (statistically insignificant)

Mutations with $\beta_1 > 0$ and $p < 0.05$ were designated as experiencing directional (positive) selection, reflecting a statistically significant increase in allele frequency over time. Mutations with $\beta_1 < 0$ and $p < 0.05$ were classified as under purifying selection, exhibiting a significant decline in frequency. All other mutations, including $p > 0.05$ or those observed in fewer than ten generations, were attributed to genetic drift, since their allele frequency changes lacked statistical support for selective trends or were insufficient in duration for inference (Natural Selection, Genetic Drift, and...; Neutral Theory: The Null Hypothesis o...).

**Supplementary Note 3. kGain distribution with population-specific evolved genomes.**

We calculated k-mer frequencies independently for each population by introducing the mutation observed at generation 57.5k into the wild-type reference genome and analyzing the resulting evolved genome. Then, for the 36,922 mutations observed in the *E. coli* LTTE experiment, we computed the kGain score based on these population-specific frequencies. We found that the kGain distribution shifts toward higher values when using population-specific evolved genomes instead of a single wild-type genome. Additionally, the median kGain increases across generations when using evolved genomes, further supporting our hypothesis. We selected generation 57.5k as the final evolved genome instead of 60k due to missing allele count data within the LTTE dataset. Generations beyond 57.5k were systematically excluded, as missing information rendered them unsuitable for analysis.

**Supplementary Note 4. Permutation test for kGain level changes with time in LTEE**

To verify the statistical significance of time series, we employed a non-parametric permutation test based on Kendall's Tau correlation to assess the association between the median of score across levels of generation number. The observed correlation was computed between the ranks of the generation number and the generation-wise median values of the score. This observed correlation was then compared against a null distribution generated from 10000 permutations, where the metric values were randomly permuted while preserving the generation-wise mutation counts. For each permutation, the generation-wise median of the permuted score within each level of the generation number was computed, and Kendall's Tau correlation was calculated between these permuted generation-wise medians and the ranks of the generation number. The Monte Carlo *P*-value was then calculated as the proportion of permutations where the permuted correlation was greater than or equal to the observed correlation using the formula:

$$\text{Monte Carlo } P\text{-value} = \frac{\sum_{i=1}^{N} \left[\tau_i \geq \tau_{obs}\right] + 1}{N + 1}$$

where $\tau_{obs}$ is the observed Kendall's Tau correlation, $\tau_i$ is the correlation from the i$^{th}$ permutation, and $N$ is the number of permutations.

**Supplementary Note 5. Modeling kGain using variant-associated mono/bi/tri nucleotide frequencies**

We evaluated whether the kGain score could be modeled using mono-, di-, and tri-nucleotide (k-mer) frequencies within variant-associated 19-mer sequences (comprising the variant itself flanked by 10 nucleotides on each side). Surprisingly, kGain, which spans genomic locations, can be predicted from local variant contexts. Regression analysis (**Supplementary Fig. S2b-g**) shows shifting k-mer associations: GC flips from negative to positive, TA from positive to negative, and tri-mers like CTA and TAG from positive to negative, while CAG and CTG shift from negative to positive. TAG, a stop codon, exemplifies this: in the reference sequence, it may support function (e.g., translation termination), but when introduced by mutation, it shows weaker kGain association. This suggests mutations adding stop codons like TAG may disrupt protein synthesis without clear selective benefit, unlike reference TAGs potentially tied to splicing or other roles.

**Supplementary Note 6. Fixed vs not fixed mutations in *E. coli* LTEE dataset**

To determine whether a mutation was fixed in a population, we examined its allele frequency during the most recent generations. A mutation that is fixed should appear in nearly all individuals, leading to a very high allele frequency at its genomic position. We assessed fixation status by checking whether the mutation maintained a high allele frequency when it was last observed. For each mutation, we specifically looked at the last two generations in which it appeared. If the allele frequency in both of these generations was greater than or equal to a predefined threshold (0.95 in our analysis), we classified the mutation as “fixed”. This means the mutation was present in almost every individual in the population by the end of the experiment. If either of the two most recent allele frequencies fell below the threshold, the mutation was considered “not fixed”. We selected the last two generations and a 0.95 threshold as criteria for this classification, but both can be adjusted or optimized in future studies as needed. This method offers a transparent way to determine fixation status, using only allele frequency information from the final generations of observation.

**Supplementary Note 7. In-house adaptation experiment control**

A control WT lineage was propagated for 7 passages without antibiotics, and whole-genome sequencing (WGS) revealed that the founder mutations persisted, plus one spontaneous mutation in *tldD*, indicating that the extensive mutations observed in D, R1, and R3 were driven primarily by antibiotic selection.

**Supplementary Note 8. SARs-CoV2 kGain insights.**

The evolution of SARS-CoV-2 exemplifies how the host-pathogen interactions shape the spontaneous emergence of mutations and new lineages that exhibit higher fitness during replication. Notably, the ORF3a gene is recognized for its diversity and functional adaptation throughout viral evolution. Specific variants of ORF3a, including ORF3a:S26L, ORF3a:T223I, ORF3a:S171L, ORF3a:Q57H, and ORF3a:S253P, have been identified as integral to the overall fitness of the virus in a study by Zhang et al. (2022). By using kGain scores, we can infer which of these variants might be favored during viral evolution.

Our analysis reveals that accessory gene mutations such as ORF3a:S26L, ORF3a:T223I, and ORF3a:S171L are associated with higher kGain scores compared to ORF3a:Q57H and

ORF3a:S253P. The former mutations are located in highly conserved regions, such as β-sheets or the S1-S2 domain, where deleterious mutations are typically omitted from the population, and beneficial ones confer survival advantages. In contrast, ORF3a:Q57H and ORF3a:S253P are situated near the C- and N-termini, which are less conserved and thus more tolerant to mutations that do not significantly impact protein function. This differential conservation likely explains the higher kGain scores observed for ORF3a:S26L, ORF3a:T223I, and ORF3a:S171L. Of relevant interest is the D614G mutation within the Spike protein of SARS-CoV-2 that was found to enhance infectivity and transmissibility relative to the original strain largely due to increased affinity for host cell receptors. The higher kGain scores associated with the D614G mutation possibly explain the influence on viral behavior.

**Supplementary Note 9. Effect size calculations**

In this study, we employ a median-based effect size instead of the traditional Cohen's *d* to enhance robustness against outliers and non-normal distributions. By using the median in place of the mean and the median absolute deviation (MAD) instead of the standard deviation, this approach provides a more stable and reliable measure of differences between distributions, particularly in cases of skewed or heavy-tailed data.

**Supplementary Figures**

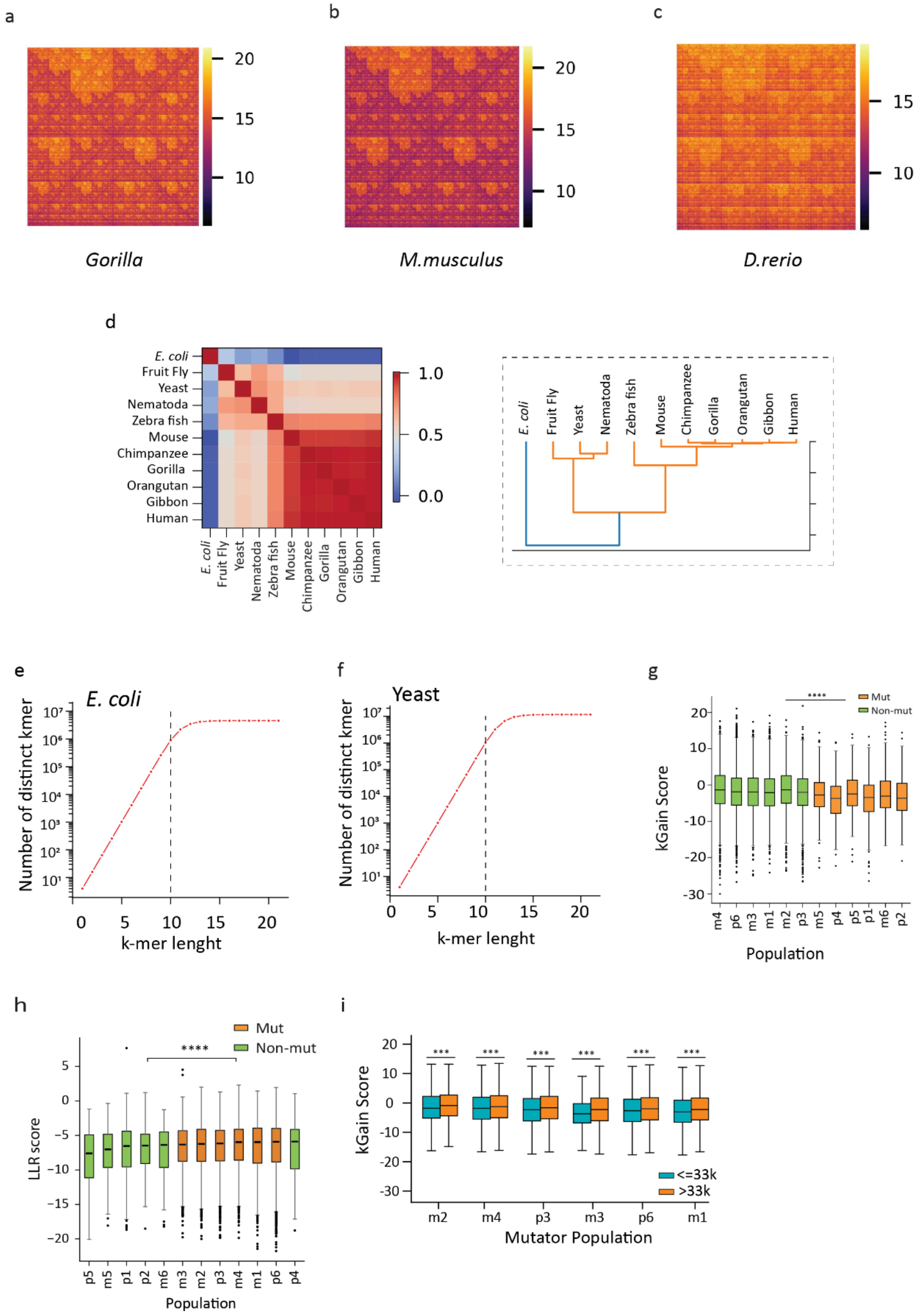

a
b
c
Gorilla
M.musculus
D.rerio
d
E. coli
Fruit Fly
Yeast
Nematoda
Zebra fish
Mouse
Chimpanzee
Gorilla
Orangutan
Gibbon
Human
e
E. coli
Number of distinct kmer
k-mer lenght
f
Yeast
Number of distinct kmer
k-mer lenght
g
Mut
Non-mut
kGain Score
Population
h
Mut
Non-mut
LLR score
Population
i
kGain Score
<=33k
>33k
Mutator Population


S1

**Supplementary Figure 1. Fitness, kGain score and LLR score across mutators and non-mutators** **(a-c)** Heatmaps illustrating normalised frequency values (in negative logarithmic scale) to reduce the range, which demonstrates that various organisms (*Gorilla*, *M. musculus*, *and D. rerio*) exhibit distinct patterns of k-mer abundance within their DNA sequences. **(d)** Heatmap of k-mer frequency correlation across species **(e-f)** Lineplot depicting variation in the number of distinct k-mers with k-mer length. **(g-h)** kGain (*P*-value between mutator vs. non-mutator is 5.42e-39) and LLR (*P*-value between mutator vs. non-mutator is 1.56e-09) scores box plot distribution for mutator and non-mutator. **(i)** The box plot shows the kGain scores in the mutator population, divided at the 33k generation mark, which is when citrate utilisation was first observed in the population (*P*-value = 4.39e-129 and effect size = 1.57e-01 for p3, *P*-value = 2.16e-201 and effect size = 1.73e-01 for p6, *P*-value = 2.25e-68 and effect size = 2.14e-01 for m1, *P*-value = 1.79e-124 and effect size = 2.37e-01 for m2, *P*-value = 5.48e-100 and effect size = 3.95e-01 for m3, *P*-value = 2.34e-50 and effect size = 1.47e-01 for m4).

[Note: The P-value cutoff for all the plots is 0.05. *, **, ***, and **** refer to P-values <0.05, <0.01, <0.001, and <0.0001, respectively]

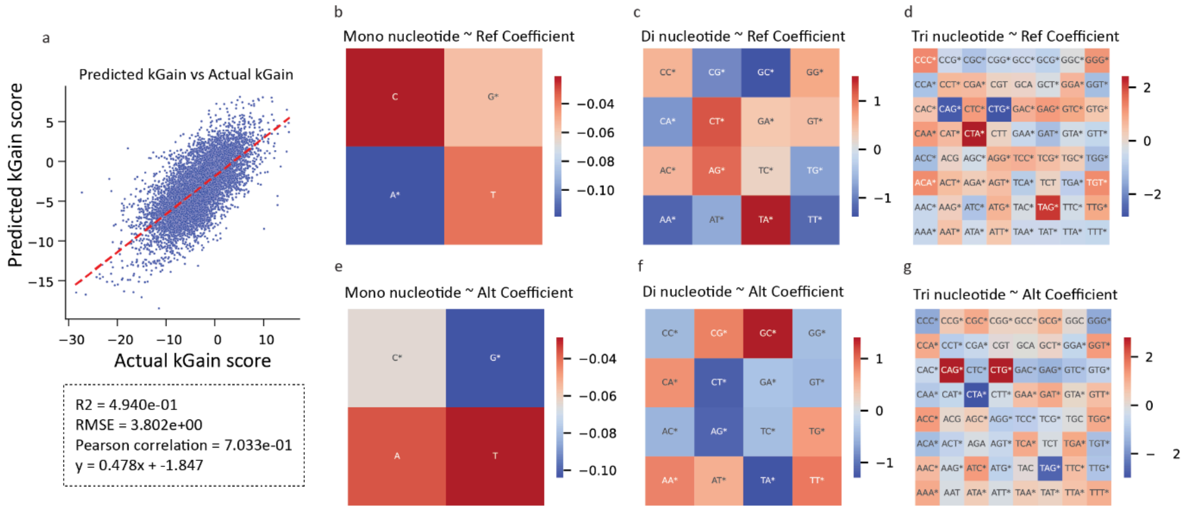
a
Predicted kGain vs Actual kGain
Predicted kGain score
Actual kGain score
R2 = 4.940e-01
RMSE = 3.802e+00
Pearson correlation = 7.033e-01
y = 0.478x + -1.847
b
Mono nucleotide ~ Ref Coefficient
c
Di nucleotide ~ Ref Coefficient
d
Tri nucleotide ~ Ref Coefficient
e
Mono nucleotide ~ Alt Coefficient
f
Di nucleotide ~ Alt Coefficient
g
Tri nucleotide ~ Alt Coefficient

S2

**Supplementary Figure 2. Prediction of the kGain score using a Ordinary Least Squares regression model. (a)** A scatter plot comparing predicted and actual kGain scores, featuring an Ordinary Least Squares regression line in red (R2 = 4.94e-01, RMSE = 3.80e+00, Pearson correlation = 7.03e-01, slope = 4.78e-01) **(b-g)** Linear regression coefficients for each mono, di, and tri-length kmer frequency from both reference and alternate sequences ( * indicate statistically significant).

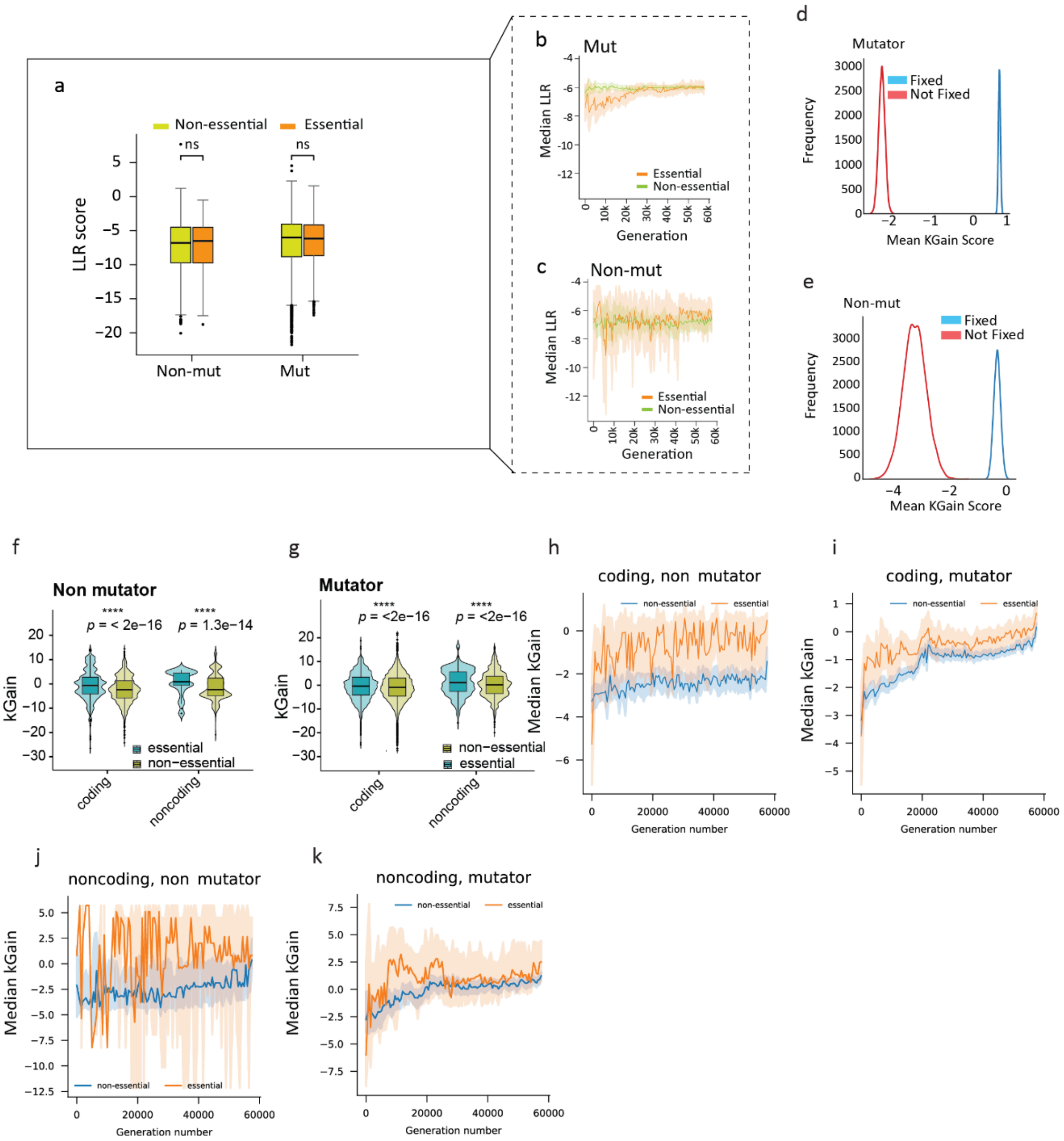
a
Non-essential
Essential
ns
ns
LLR score
5
0
−5
−10
−15
−20
Non-mut
Mut
b
Mut
Median LLR
Essential
Non-essential
Generation
c
Non-mut
Median LLR
Essential
Non-essential
Generation
d
Mutator
Fixed
Not Fixed
Frequency
Mean KGain Score
e
Non-mut
Fixed
Not Fixed
Frequency
Mean KGain Score
f
Non mutator
p = < 2e−16
p = 1.3e−14
kGain
essential
non−essential
coding
noncoding
g
Mutator
p = <2e−16
p = <2e−16
kGain
non−essential
essential
coding
noncoding
h
coding, non mutator
Median kGain
Generation number
i
coding, mutator
Median kGain
Generation number
j
noncoding, non mutator
Median kGain
Generation number
k
noncoding, mutator
Median kGain
Generation number

**Supplementary Figure 3. Fitness, kGain score and LLR score across mutators and non-mutators**

**(a)** Boxplot to demonstrate the distribution of LLR scores among mutator and non-mutator groups for essential and non-essential genes (*P*-values are 5.06e-01 for mutator and 3.86e-01 for non-mutator). **(b-c)** Generation-wise median LLR scores across mutator and non-mutator populations for essential and non-essential gene distribution. **(d)** Bootstrapped distributions of mean kGain scores for fixed and non-fixed mutations in mutator populations, generated using 10,000 iterations and subsampling 90% of the minimum group size per iteration. **(e)** Bootstrapped distributions of mean evolved kGain scores for fixed and non-fixed mutations in non-mutator populations, generated using 10,000 iterations and subsampling 90% of the minimum group size per iteration. **(f-g)** Violin and box plots showing the distribution of kGain scores for essential and non-essential genes in coding and noncoding regions, separated by non-mutator (*P*-value = 1.58e-05 and effect size = 3.31e-01 for coding, *P*-value = 6.50e-01 and effect size = -2.55e-02 for noncoding) and mutator populations (*P*-value = 3.87e-02 and effect size = 7.38e-02 for coding, *P*-value = 1.42e-01 and effect size = 7.99e-02 for noncoding). **(h-i)** Trajectories of median kGain scores over generations for coding regions, comparing essential and non-essential genes in non-mutator and mutator populations. **(j-k)** Median kGain trajectories for noncoding regions over generations in non-mutator and mutator populations, plotted separately for essential and non-essential genes.

[Note: The P-value cutoff for all the plots is 0.05. *, **, ***, and **** refer to P-values <0.05, <0.01, <0.001, and <0.0001, respectively]

a

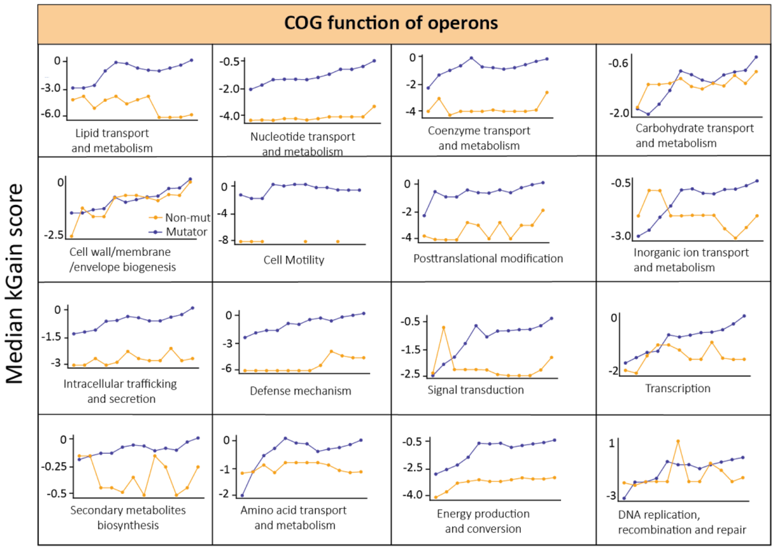

COG function of operons
Median kGain score
Lipid transport and metabolism
Nucleotide transport and metabolism
Coenzyme transport and metabolism
Carbohydrate transport and metabolism
Cell wall/membrane /envelope biogenesis
Non-mut
Mutator
Cell Motility
Posttranslational modification
Inorganic ion transport and metabolism
Intracellular trafficking and secretion
Defense mechanism
Signal transduction
Transcription
Secondary metabolites biosynthesis
Amino acid transport and metabolism
Energy production and conversion
DNA replication, recombination and repair


S4

**Supplementary Figure 4. Dynamics of median kGain scores across functional categories of Operons.**

Line plots show the median kGain score over generations for Operons grouped by COG functional categories, comparing mutator (blue) and non-mutator (orange) populations. Each panel represents a different COG function, illustrating distinct evolutionary trends in metabolic, regulatory, and cellular processes between population types.

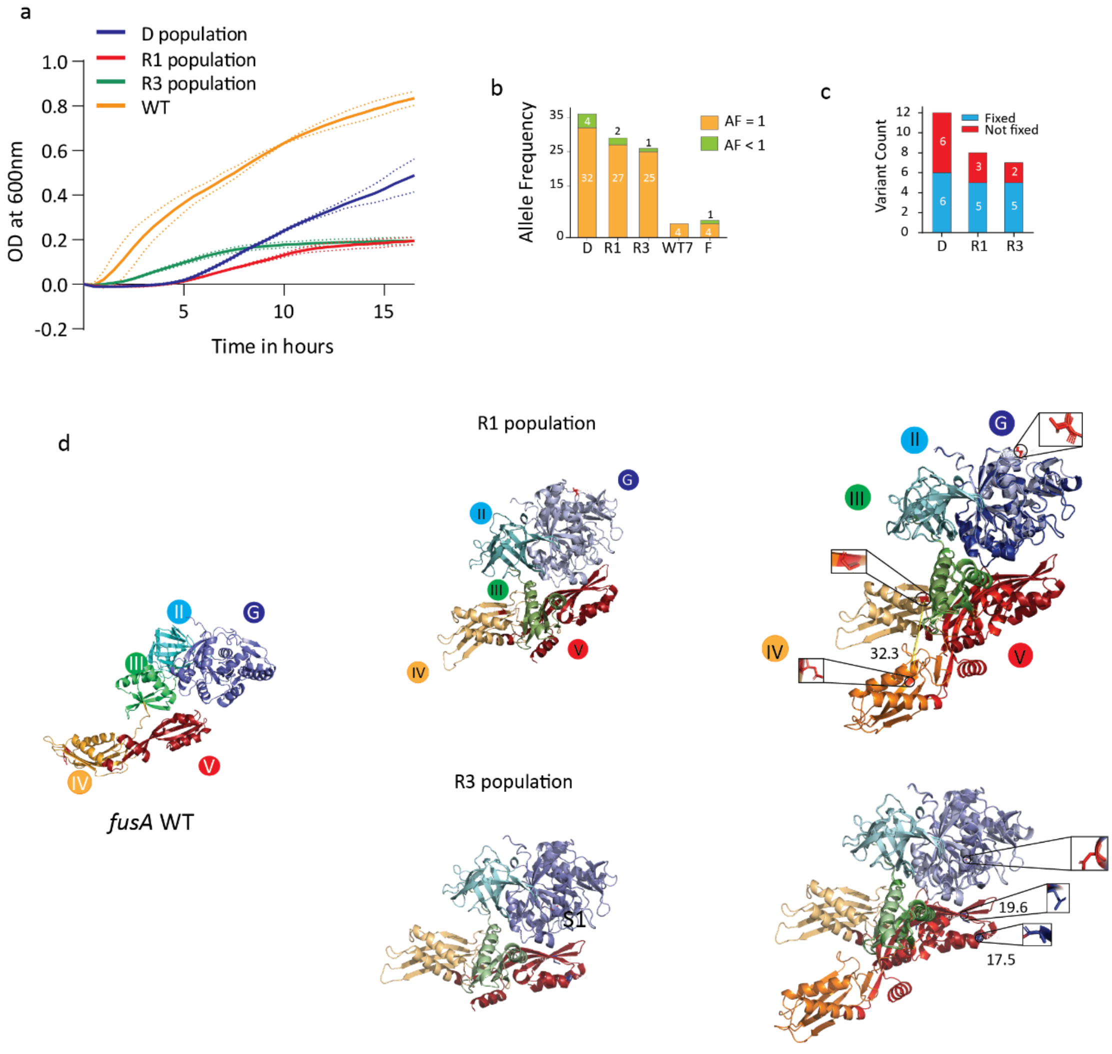
a
D population
R1 population
R3 population
WT
OD at 600nm
Time in hours
b
Allele Frequency
AF = 1
AF < 1
D R1 R3 WT7 F
c
Variant Count
Fixed
Not fixed
D R1 R3
d
R1 population
R3 population
fusA WT
32.3
19.6
17.5

S5

**Supplementary Figure 5.** ***E.coli*** **adaptation experiment**

**(a)** Growth curve for passage 5 of the D, R1, R3 populations, and the WT in LB medium with antibiotics. **(b)** Mutation count across populations (D, R1, R3, WT7, and F) across allele frequency categories. **(c)** Mutation count across populations (D, R1, R3, WT7, and F), comparing the fixed (blue) and non-fixed (red) populations across generations. **(d)** Structural alignment of WT fusA and its mutant variants from R1 and R3 populations, highlighting differential folding patterns between the WT and mutant proteins.

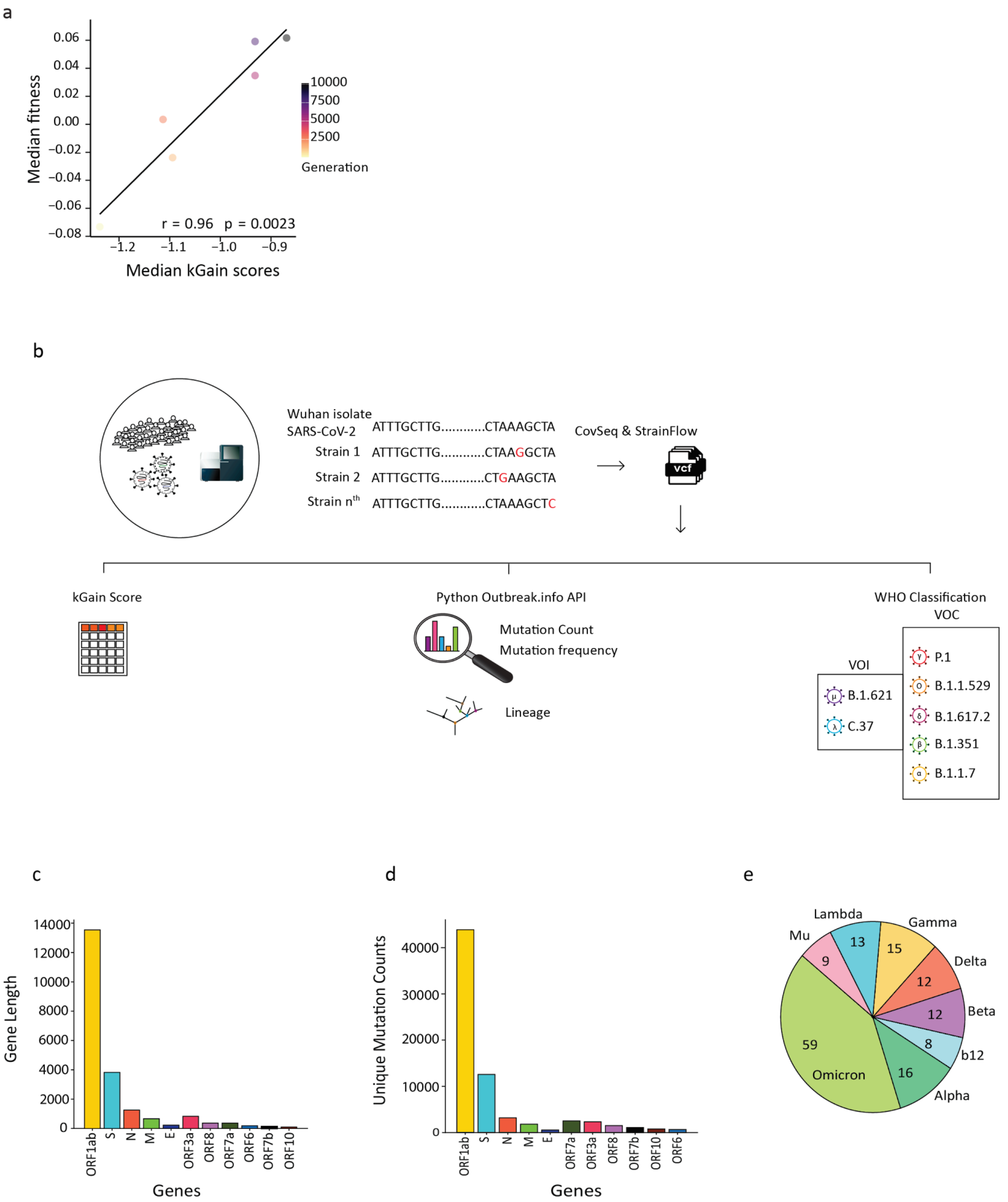
a
Median fitness
Median kGain scores
r = 0.96 p = 0.0023
Generation
10000
7500
5000
2500
b
Wuhan isolate SARS-CoV-2 ATTTGCTTG...........CTAAAGCTA
Strain 1 ATTTGCTTG...........CTAAGGCTA
Strain 2 ATTTGCTTG...........CTGAAGCTA
Strain nth ATTTGCTTG...........CTAAAGCTC
CovSeq & StrainFlow
vcf
kGain Score
Python Outbreak.info API
Mutation Count
Mutation frequency
Lineage
WHO Classification
VOC
VOI
B.1.621
C.37
P.1
B.1.1.529
B.1.617.2
B.1.351
B.1.1.7
c
Gene Length
Genes
ORF1ab
S
N
M
E
ORF3a
ORF8
ORF7a
ORF6
ORF7b
ORF10
d
Unique Mutation Counts
Genes
e
Lambda 13
Gamma 15
Mu 9
Delta 12
Beta 12
b12 8
Alpha 16
Omicron 59

S6

**Supplementary Figure 6. Yeast and *SARS-CoV-2* Analysis**

**(a)** For every available generation, we computed the median kGain and median fitness across the entire population. These values were then visualised in the form of a scatter plot, where the color gradient represents generation in yeast. **(b)** Overview of *SARS-CoV-2* analysis pipeline. **(c)** Bar plot showing the length of genes of the *SARS-CoV-2* variant. **(d)** Bar plot showing unique mutation counts in each gene. **(e)** Pie chart illustrating the distribution of variant counts in each SARS-CoV-2 lineage.

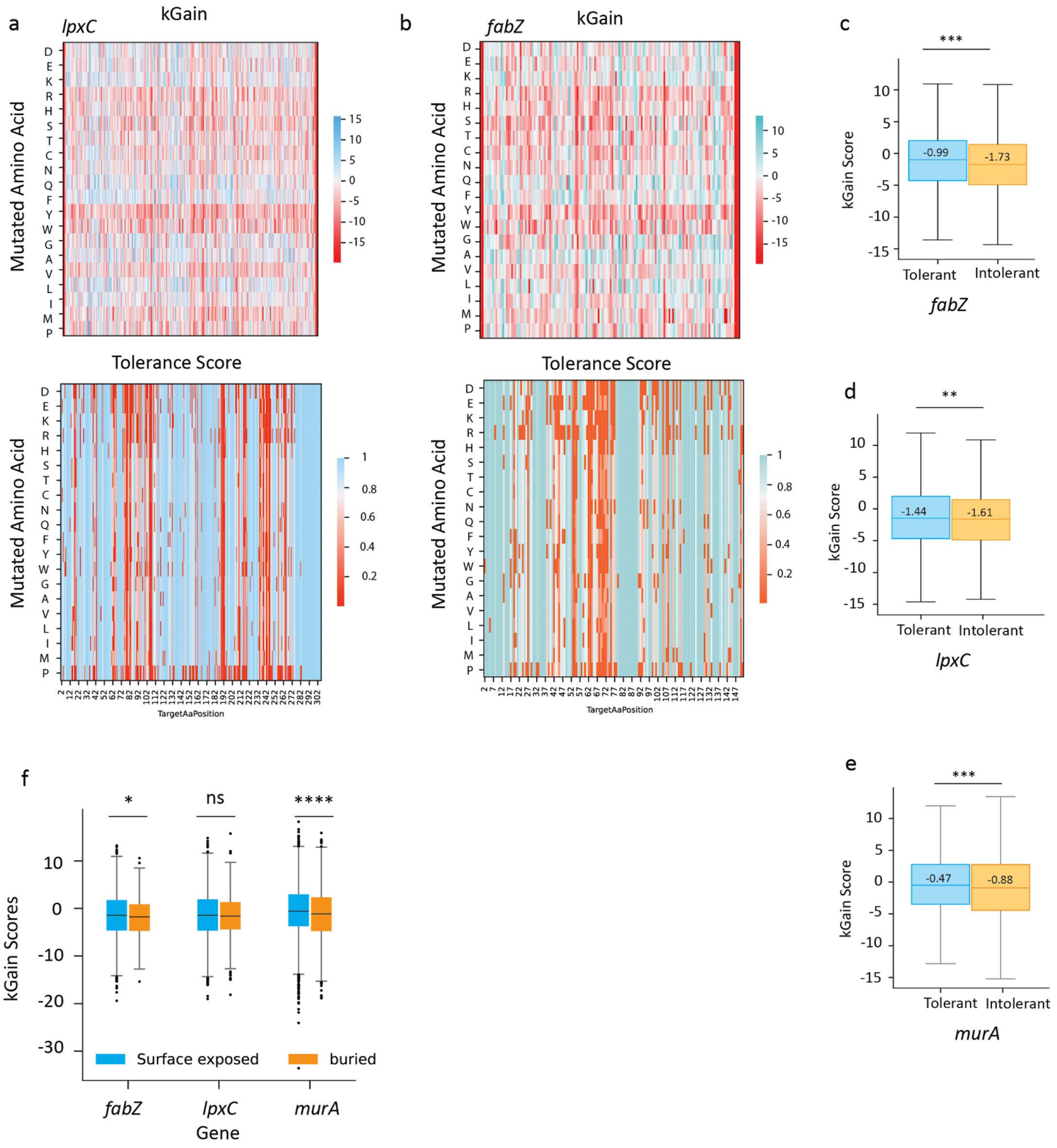
a
lpxC
kGain
Mutated Amino Acid
D E K R H S T C N Q F Y W G A V L I M P
15 10 5 0 -5 -10 -15
Tolerance Score
1 0.8 0.6 0.4 0.2
TargetAaPosition
b
fabZ
kGain
Mutated Amino Acid
10 5 0 -5 -10 -15
Tolerance Score
1 0.8 0.6 0.4 0.2
TargetAaPosition
c
***
kGain Score
-0.99
-1.73
Tolerant
Intolerant
fabZ
d
**
kGain Score
-1.44
-1.61
Tolerant
Intolerant
lpxC
e
***
kGain Score
-0.47
-0.88
Tolerant
Intolerant
murA
f
*
ns
****
kGain Scores
Surface exposed
buried
fabZ
lpxC
murA
Gene

S7

;

**Supplementary figure 7. kGain scores dynamics captured in DMS studies**

**(a-b)** Heatmaps displaying kGain scores for mutations across different amino acids in the *lpxC* and *fabZ* genes. The color scale represents the range of kGain scores, with red indicating negative values and sky blue representing positive values. Corresponding heatmaps illustrate tolerance scores, with the color scale ranging from red to blue, representing tolerance values from 0 to 1. **(c-e)** Boxplots comparing kGain scores for *fabZ (P*-value: 3.69e-4*), lpxC (P*-value: 3.48e-3*),* and *murA (P*-value: 1.60e-4*)* mutations, categorised by tolerance levels. Tolerant mutations (Blue) are associated with higher kGain scores, while intolerant mutations (Orange) show significantly lower values. **(f)** Boxplot depicting Relative Solvent Accessibility (RSA) scores across mutations. The plot shows the distribution of kGain scores in relation to RSA values, highlighting differences between buried and surface-exposed residues (*P*-values are 1.79e-02 for fabZ, 2.22e-01 for lpxC , and 1.87e-05 for murA).

[**Note**: The p-value cutoff for all the plots is 0.05. *, **, ***, and **** refers to p-values <0.05, <0.01, <0.001, and <0.0001, respectively.]

| Genes | Gene group | Unique SNPs | Unique indels |
|---|---|---|---|
| S (Spike) | Structural | 8716 | 3866 |
| E (Envelope) | Structural | 491 | 72 |
| M (Membrane) | Structural | 1553 | 264 |
| N (Nucleocapsid) | Structural | 2707 | 496 |
| ORF1ab | Non-structural | 38319 | 5529 |
| ORF3a, ORF6, ORF7a, ORF7b, ORF8, ORF10 | Accessory | 5185 | 3746 |

**Table S1. Data dictionary mapping gene, gene group, no of unique SNPs and indels in *SARS-CoV-2***

| PANGO lineage | WHO classification | Variant classification |
|---|---|---|
| B.1.1.7 | Alpha | VOC |
| B.1.351 | Beta | VOC |
| P.1 | Gamma | VOC |
| B.1.617.2 | Delta | VOC |
| BA.1, BA.2, BA.3, BA.5, B.1.1.529 | Omicron | VOC |
| C.37 | Lambda | VOI |
| B.1.621 | Mu | VOI |
| B.1.2 | - | VOI |

**Table S2. The dataset comprises information pertaining to PANGO lineages, their WHO and variant classification in *SARS-CoV-2***
